# Activated fibroblasts drive cellular interactions in end-stage pediatric hypertrophic cardiomyopathy

**DOI:** 10.1101/2024.01.25.577226

**Authors:** Hanna J. Tadros, Diwakar Turaga, Yi Zhao, Chang-Ru Tsai, Iki A. Adachi, Xiao Li, James F. Martin

## Abstract

Hypertrophic cardiomyopathy (HCM) is a relatively rare but debilitating diagnosis in the pediatric population and patients with end-stage HCM require heart transplantation. In this study, we performed single-nucleus RNA sequencing on pediatric HCM and control myocardium. We identified distinct underling cellular processes in pediatric, end-stage HCM in cardiomyocytes, fibroblasts, endothelial cells, and myeloid cells, compared to controls. Pediatric HCM was enriched in cardiomyocytes exhibiting “stressed” myocardium gene signatures and underlying pathways associated with cardiac hypertrophy. Cardiac fibroblasts exhibited clear activation signatures and heightened downstream processes associated with fibrosis, more so than adult counterparts. There was notable depletion of tissue-resident macrophages, and increased vascular remodeling in endothelial cells. Our analysis provides the first single nuclei analysis focused on end-stage pediatric HCM.

## Introduction

Hypertrophic cardiomyopathy (HCM) is a relatively rare, but debilitating diagnosis in the pediatric population, with an estimated incidence of 2.9/100,000 [1]. The outcomes and presentations depend on underlying etiology and age of presentation and the phenotype ranges from mild cardiac hypertrophy to sudden cardiac death and end-stage heart failure [2]. In fact, young HCM patients carry an almost 4-fold higher mortality compared to the general population [3]. The “cumulative burden” of HCM across the lifetime of a patient is significant and younger patients carry a heightened risk of adverse outcomes. In particular, end-stage pediatric HCM patients represent a vulnerable population; approximately a quarter of these patients experience significant arrhythmias and waitlist mortalities can be as high as 33% in the infant population [4].

Alterations in the structure and function of the sarcomere is thought to be the primary cause of HCM, with most variants identified in myosin heavy chain 7 encoding-*MYH7* and myosin binding protein C3 encoding-*MYBPC3* [5]. Variants generally lead to functional protein changes like reduction of ATP-responding, sensitivity of actin-myosin complex disassociation, and efficiency of force generation [5],[6]. This leads to downstream molecular changes including activation of signaling pathways, like transforming growth factor beta and mitogen-activated protein kinase pathways [5]. Ultimately, these alterations lead to histological changes like myocyte disarray, cardiac hypertrophy, and interstitial fibrosis [5].

Single-nucleus RNA sequencing (snRNA-seq) is a powerful tool used to assess gene expression across different cell types and conditions in the heart. Recently, studies utilizing snRNA-seq have described the transcriptome in adult HCM patients [7],[8]. These studies establish enrichment of genes associated with “cardiac stress” in cardiomyocytes, genes associated with fibroblast activation and fibrosis in fibroblasts, and macrophage activation and subtype switching [7],[8]. In addition, enhanced intercellular communication, both in number and strength of interactions, is seen in HCM, similar to other cardiac diseases like congenital heart disease [8],[9]. snRNA-seq studies involving cardiac disease in pediatric patients are scarce, owing to the rarity of pediatric samples [9],[10]. There have been no studies focusing on pediatric HCM.

We set out to perform snRNA-seq to study transcriptome differences between end-stage pediatric HCM patients and controls without structural heart disease. We also set out to identify pathways implicated in each of the major cell types - cardiomyocytes, fibroblasts, endothelial cells, and myeloid cells - determine co-expressed gene networks upregulated in disease, predict intercellular communications across each of the major cell types, and compare identified major pediatric HCM cellular processes to adult HCM counterparts.

## Methods

### Research ethics for donated tissues

Cardiac tissue samples used in this study were collected during cardiothoracic surgery performed at Texas Children’s Hospital (Houston, Texas). The protocols for the procurement and use of these patient sample were approved by the Institutional Review Board for Baylor College of Medicine and Affiliated Hospitals (Protocol Number H-26502). With the help of the Heart Center Biorepository at Texas Children’s Hospital, consent was obtained from patients.

### Clinical cohort

We identified four, non-syndromic, end-stage pediatric (<18 years) HCM patients transplanted at Texas Children’s Hospital/Baylor College of Medicine from 2014-2021. Of the four, we obtained left ventricular tissue samples from three at the time of heart transplant (HT; Figure 1A/1B). All three patients met indications for transplantation due to severe diastolic dysfunction leading to restrictive physiology and associated symptoms and had variants identified in sarcomere genes *TNNT2* or *MYH7*. Data for our HCM cohort was collected retrospectively via chart review from medical records. Demographics and clinical variables collected included sex, age of diagnosis, age at HT, pre-HT medications, gene sequencing testing, and laboratory findings, specifically B-type natriuretic peptide (BNP) levels. Diagnostic imaging studies collected included 2-dimensional echocardiographic studies, performed on commercially available cardiac ultrasound scanners.

**Figure 1:**
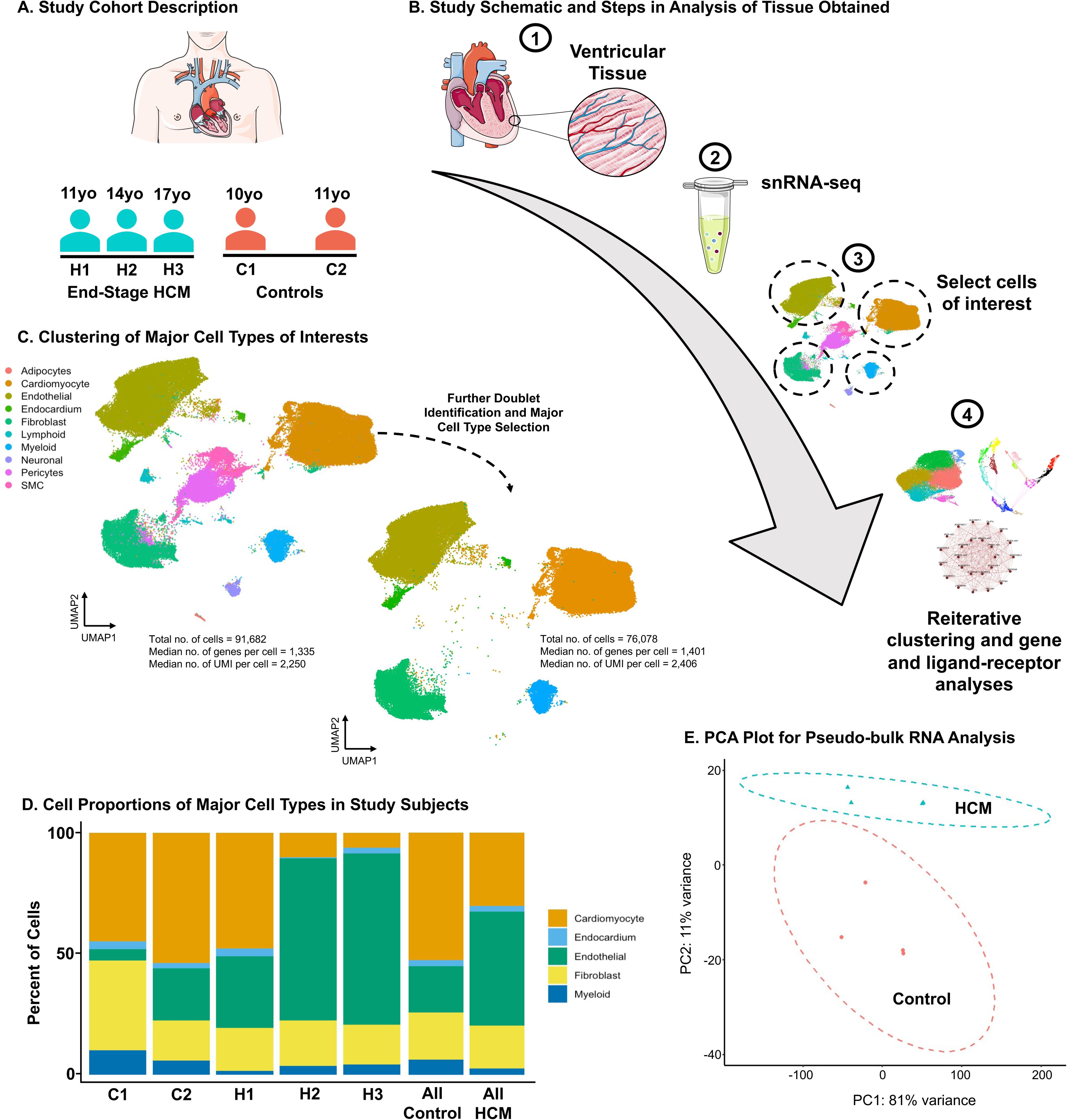
Single-nuclei analysis in end-stage, pediatric hypertrophic cardiomyopathy. A) Study cohort description exhibiting HCM and control subjects with their respective ages. B) Study schematic and steps in analysis in both controls and those with HCM from time of procurement (1), tissue handling and RNA sequencing (2), identification of major cell types of interest (3), and downstream analyses (4). C) UMAP embedding of major cell types and subsequent clustering of major cell types of interest. D) Proportion of major cell types of interest within control subjects (C1 and C2) and HCM subjects (H1, H2, and H3) and proportions in pooled cohorts (Control vs HCM). E) PCA plot for pseudo-bulk RNA-seq analysis for all major cell types of interest by diagnosis (red: Control; blue: HCM). Ellipses represent the 95% confidence intervals.

### Sample collection and nuclear Isolation

HCM tissues was collected in the operating room during HT. The anatomic location of tissue collected is LV free wall. Cardiac tissue sample was kept in cold saline on ice during transfer to the laboratory for preservation. Cardiac tissue samples were carefully dissected into multiple aliquots, which were flash-frozen and stored at –80 °C. Nuclear isolation was performed as described previously [9],[11]. Briefly, frozen cardiac tissue was dissociated by using a Dounce homogenizer. Single nuclei were isolated via fluorescence-activated cell sorting (FACS).

### Single-nucleus RNA sequencing

SnRNA-seq was performed by using the 10X Genomics platform. Isolated nuclei were loaded into the 10X Genomics Chromium Controller to obtain the gel beads in emulsion. The sequencing libraries were then prepared according to the manufacturer’s protocols for the Single-cell 3’ Reagents Kits v3. Sequencing was performed by using the NovaSeq 6000 system.

### Controls

Since our patient cohort was made up of female patients, we identified two female controls with relatively similar ages. Clinical characteristics are described in Table S1. Controls tissue data were obtained from the following published data sets [9],[10],[12]. Control data were downloaded in the format of raw sequencing reads and were re-processed with the HCM data using the same pipeline.

### SnRNA-seq data processing and integration

All newly generated and published snRNA-seq data sets were processed using a uniformed pipeline. Raw sequencing reads were aligned to the genome (build GRCh38) using the 10X Genomics toolkit CellRanger version 5.0.1 (cellranger count) with --include-introns set to true. All other parameters were left as defaults. Quality control metrics generated by CellRanger were inspected for each library. To remove background signals from ambient transcripts, the raw UMI count matrices were further processed by CellBender version 0.1.0 (cellbender remove-background) with --total-droplets-included = 25000, --low-count-threshold = 15, and --epochs = 200. To minimize the loss of valid cell barcodes called by CellRanger, we also set --expected-cells at 1.5 times of CellRanger output nuclei number. The output matrices from CellBender were filtered to only include valid cell barcodes that were also identified by CellRanger. Additional quality controls at single nucleus level were performed for each library. Briefly, we first identified low-quality nuclei based on fixed cut-offs of UMI count per nucleus > 200, gene count per nucleus > 150 and mitochondria gene-derived UMI <5%. Then, a set of dynamic cut-offs based on per-library data distribution were calculated, which is essential to account for heterogeneity between samples. In brief, for each library, an upper and lower bound were set at the 75th percentile plus 1.5 times the interquartile range (IQR) and the 25th percentile minus 1.5 times IQR, respectively, for UMI count and gene count per nuclei. Next, the remaining nuclei were evaluated by the Scrublet tool [13] to identify potential doublets, with parameters expected_doublet_rate = 0.1 and call_doublets threshold = 0.25. Finally, we integrated all samples and corrected batch-effect using a deep generative model scANVI [14]. The scANVI latent space was reduced to generate the final global UMAP embedding and subsequent subcluster UMAP embeddings for cardiomyocytes, fibroblasts, endothelial/endocardial cells, and myeloid cells.

### Sublustering and differential gene expression testing

We subsetted data by major cell type, focusing on cardiomyocytes, fibroblasts, endothelial/endocardial cells, and myeloid cells. Re-clustering was then performed with the FindNeighbors function using 50 dimensions of the scVI latent space and subsequently the FindCluster function with a Louvain resolution of 0.2. Clusters expressing markers of two major cell types were labeled as doublets and removed. Testing of differentially expressed genes (DEGs) between clusters was completed using the FindAllMarkers function (Wilcoxon rank-sum test, min. pct = 0.05, thresh.use = 0.15). This function employs the Bonferroni correction to calculate an adjusted *P* value. We next sorted DEGs by the log fold-change of the average expression (avg log2FC or FC) and filtered for DEGs with adjusted *P* values <0.05. Sample level pseudobulk differential expression analysis was performed after extracting the single-cell RNA-seq raw count data and using the typical workflow with the DEseq2 package (version 1.38.3; https://hbctraining.github.io/scRNA-seq/lessons/pseudobulk_DESeq2_scrnaseq.html) [15]. To explore the similarity of the samples based on the condition of interest (HCM vs control), we performed and plotted the principal component analysis (PCA).

### Differential abundance testing

For differential abundance testing, we used the MiloR package (version 1.6.0) to determine the abundance of diseased cohort control neighborhoods in clusters.[16] For k-nearest graph construction, we used the reduced dimensions output from scVI and depending on the cell number in each major cell type, we set the number of nearest neighbors (k) from 10 to 50. For counting cells in neighborhoods and differential abundance testing, we inputted the samples (ie. tissue samples) used and matched them to the condition of interest (ie. HCM vs control). An alpha threshold of 0.1 for plotNhoodGraphDA was used to generate the final graph of neighborhoods. *P* values were adjusted with the Benjamini-Hochnberg correction.

### Weighted gene co-expression network analysis

To perform weighted gene co-expression network analysis (WGCNA), we used the high dimensional WGCNA (hdWGCNA) package (version 0.2.26) [17]–[19]. The recommended hdWGCNA workflow was followed. We included genes expressed in at least 5% of the cells in each dataset. To construct the metacell expression matrices, we inputted the samples as recommended in the group.by parameter to ensure that cells coming from the same biological sample of origin are used to construct the metacells. The number of cells to be aggregated (k) was determined by the number of cells in each dataset and ranged from 25-50 and the maximum number of shared cells between two metacells was set to 10. To perform k-nearest neighbors, the scVI dimensionality reduction was used. We then focused on the “hub” genes and their respective eigengene-based connectivity (kME). The top 25 “hub” genes were identified based on their kME, a measure of the correlation of a gene and the respective module eigengene, and was used to compute a gene score and module networks. We then ran the UMAP algorithm on the topological overlap matrix, set the number of hub genes to include in the UMAP embedding to 10, the neighbor’s parameter to 15, and the minimum distance between points at 0.1, and finally ran the ModuleUMAPPlot function to plot the genes and their co-expression relationships.

To perform differential module eigengene analysis, we subset the data to group cells from patients with HCM and controls separately. hdWGCNA employs a function analogous to the FindMarkers function in Seurat called the FindDMEs function, which uses the Wilcoxon rank-sum test to perform differential module eigengene expression testing. To test for similarities in our Seurat and hdWGCNA analyses, we then compare hdWGCNA modules to cluster marker genes. We reran the FindAllMarkers function with a log fold change threshold of 1 to identify highly DEGs in each of our clusters and computed marker gene overlap. In both the former analyses, *P* values were adjusted with Benjamini-Hochnberg correction and we set a significance of 0.05.

### Module score creation

We utilized the AddModuleScore function in the Seurat package to develop cell scores for myocardial stress, LAMININ ligand, COLLAGEN ligand, and fibroblast activation, based on previous genetic signatures defined by Kuppe et al., Chaffin et al., and CellChat (https://www.cellchat.org/cellchatdb/) [7],[20]. This function normalizes the average expression of a given gene set against the average expression of control genes across the whole dataset. Scores were then compared within each of the cardiomyocyte and fibroblast clusters, between HCM and control cells, and between adult and pediatric HCM. The genes included in each of these scores are listed in Table S2.

### Functional enrichment analyses and cell-cell Interactions

To test for functional enrichment, analyses were performed following DEG or “hub” gene identification in the Seurat and the hdWGCNA analyses, respectively. The clusterProfiler package (version 4.6.2) was used to perform enrichGO and enrichKEGG analyses on the top 500 genes to identify functional pathways enriched in each cluster and similarly in pediatric vs adult HCM [21]. As part of the hdWGCNA package, EnrichR enrichment testing was performed on the top 500 genes within each respective module to identify functional pathways enriched within each module. Enriched terms and pathways were filtered based on adjusted *P* values of <0.05.

Intercellular interactions were analyzed with CellChat (version 1.6.1) [22]. The typical workflow and default parameters were followed. In short, control and HCM Seurat objects were subsetted into two separate datasets and converted into CellChat objects. To compare intercellular interactions amongst major cell types in HCM and control samples, the CellChat objects were combined and the CellChat comparison analysis of multiple datasets workflow was followed. Given we saw a significantly higher number and greater weight of interactions in our HCM cohort, we then further analyzed intercellular communication in the HCM dataset alone, stratified by individual cluster.

### Statistical analysis and data visualization

All statistical analyses outside the previously mentioned packages were completed using the ggpubr (version 0.6.0) package in R (version 4.2.3). Box plots exhibit medians, the lower and upper quartiles, and the minimum and maximum data values. Relative expression between two groups were compared with the Wilcoxon rank-sum test and more than two groups were compared with the Kruskal Wallis test. Data visualization was completed via the various cited packages described above. In addition to these, bar plots depicting percentages were created using dittoSeq [23]. Pre-mutation testing between cell proportions was completed with the scProportionTest package [24]. Parts of Figure 1 were drawn by using pictures from Servier Medical Art. Servier Medical Art by Servier is licensed under a Creative Commons Attribution 3.0 Unported License (https://creativecommons.org/licenses/by/3.0/).

## Results

### Global SnRNA-seq analysis reveals differences in cellular composition in HCM

From the 8 tissue samples, we generated 91,682 single-cell transcriptomes and identified 10 clusters representing major cell types after batch correction and doublet removal (Figure 1C). Annotation of major cell types was completed based on differential expression of marker genes. We then focused on cardiomyocytes, fibroblasts, endothelial/endocardial cells, and myeloid cells. Following re-clustering, we further manually removed doublets for a total remaining 29,501 cardiomyocytes, 29,512 endocardial and endothelial cells, 13,959 fibroblasts, and 3,106 myeloid cells (Figure 1C). Proportions of total cells contributed to each major cell type by subject and by total HCM and control are exhibited in Figure 1D. Premutation testing exhibited a significantly increased proportion of endothelial cells (Log2FD= 1.29, Adjusted *P-*value=0.001) and decreased proportion cardiomyocytes (Log2FD= -0.80, Adjusted *P-*value <0.001), fibroblasts (Log2FD= -0.12, Adjusted *P-*value <0.001), and myeloid cells (Log2FD= - 1.27, Adjusted *P-*value <0.001) in HCM. Sample level pseudo-bulk RNA-seq analysis exhibited global transcriptional differences between HCM and control samples (Table S3). The influence of disease on PCA analysis is exhibited in Figure 1E and PC2 distinguishes control and HCM samples.

### Cardiomyocyte gene expression implicates several pathways enriched in HCM-enriched clusters

Unbiased re-clustering of the cardiomyocyte population identified 6 cardiomyocyte clusters denoted as CM1-CM6 (Figure 2A). Proportions of cells contributed to each cluster by subject and total HCM and control are exhibited in Figure 2B. Premutation testing exhibiting a significantly increased proportion of HCM-derived cells in CM1 (Log2FD= 2.72, Adjusted *P-*value<0.001) and CM4 (Log2FD= 1.31, Adjusted *P-* value<0.001) and increased proportion of control-derived cells in CM2 (Log2FD= 0.58, Adjusted *P-*value<0.001), CM3 (Log2FD= 2.32, Adjusted *P-*value<0.001), CM5 (Log2FD= 2.69, Adjusted *P-*value<0.001), and CM6 (Log2FD= 5.58, Adjusted *P-*value <0.001). This was further substantiated in the differential abundance analysis where the CM1 and CM4 clusters were enriched with HCM neighborhoods (Figure 2C).

**Figure 2:**
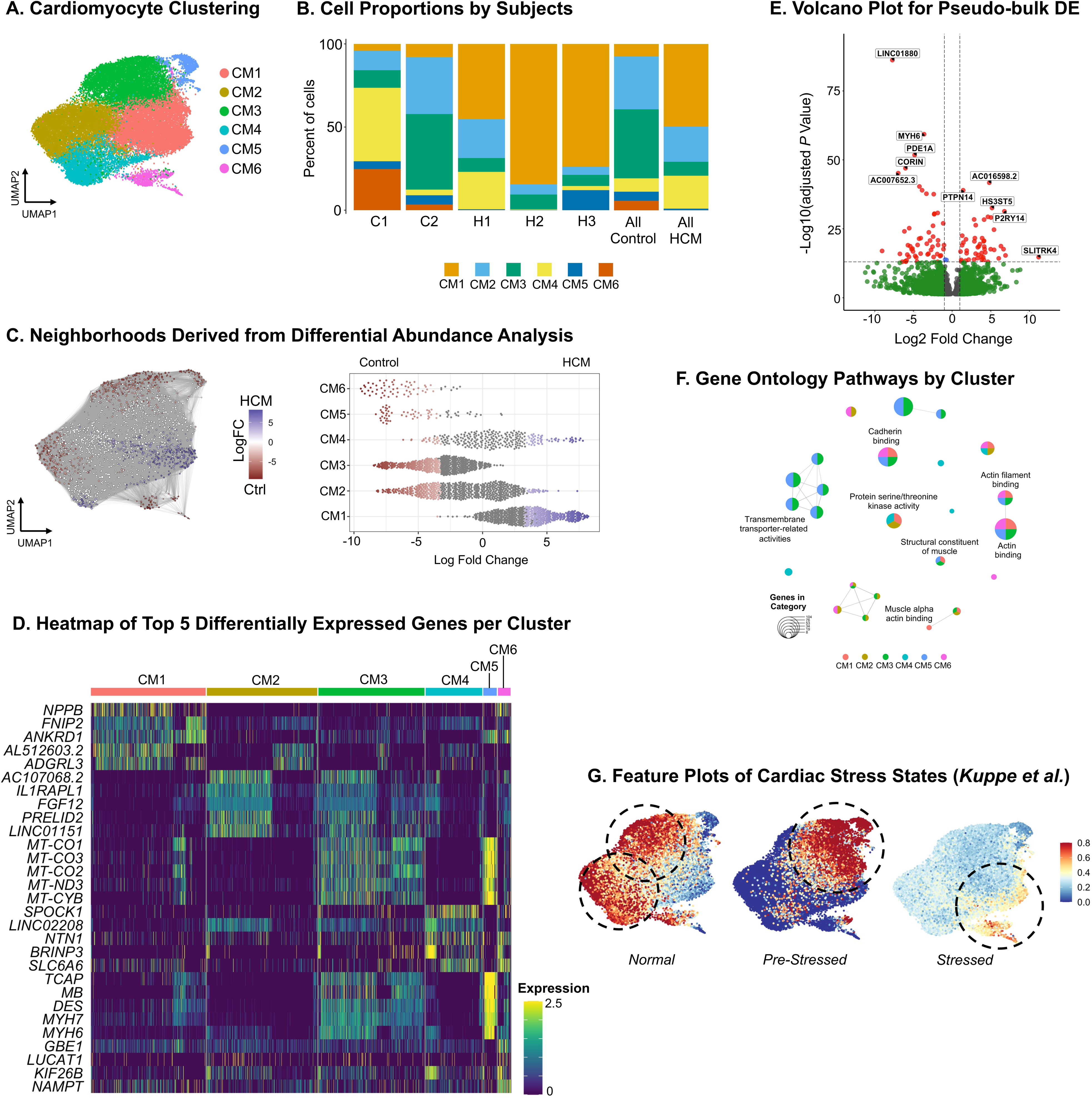
HCM cardiomyocytes are enriched in insulin receptor and actin binding pathway genes. A) UMAP embedding of reiterative clustering of cardiomyocytes. B) Bar plot depicting proportion of major cell types of interest within control subjects (C1 and C2) and HCM subjects (H1, H2, and H3) and proportions in pooled cohorts (Control vs HCM). C) Left: Embedding derived from Milo k-nearest neighborhood differential abundance testing and layout derived from UMAP embedding. Nodes with false discovery rate <10% depict neighborhoods and are colored by log fold changes for HCM (blue) versus control (red) samples. Right: Beeswarm plot depicting significant log fold changes for HCM (blue) and control (red) neighborhoods derived from differential abundance testing stratified by cardiomyocyte clusters (y-axis). D) Heatmap of top 5 differently expressed genes derived from FindAllMarkers function in CM1, CM2, CM3, CM4, CM5, and CM6. E) Volcano plot derived from pseudo-bulk RNA differential gene expression. F) Enrichment map stratified by CM cluster depicting the proportional GO pathway analysis. Node size is determined by the number of genes within each GO pathway. G) Feature plots illustrating expression of normal, pre-stressed, and stressed cardiomyocyte scores derived from Kuppe et al. in combined HCM and control cells, overlaid on the original UMAP embedding.

In the DEG analysis, DEGs that are significantly upregulated (FC >1) in CM1 have been previously implicated in cardiomyopathy and are associated with “cardiac stress” including *ANKRD1, NPPB,* and *PROS1* (Figure 2D, Table S4) [8],[25]. Other upregulated genes of interest include *PLCE1,* which promotes myocardial ischemia-reperfusion injury, and *IGF1R*, shown to be related to cardiac homeostasis and aging [26],[27]. Interestingly, control-dominant CM5 exhibited upregulation of *TCAP, MB, DES, MYH7,* and *XIRP2,* several of which are known to be upregulated in stressed myocardium, as well as other sarcomere genes like *TNNT2, TNNC1,* and *TNNI3.* Pseudobulk-RNA analysis exhibited some overlap with HCM dominant clusters in our DEG analysis. It also identified other genes of interest like upregulated *SLITRK4*, previously shown to be diagnostic marker in HCM [28]. Additionally, genes known to be downregulated in end-stage cardiomyopathy like *MYH6* and *CORIN* were identified (Figure 2E; Table S5) [9],[12]. Functional enrichment analysis revealed upregulation of Gene-Ontology (GO) pathways related to actin binding in HCM-enriched CM1, but were also seen in control-enriched CM3, CM5, and CM6 (Figure 2F, Table S6). Upregulation of insulin receptor pathways was unique to CM1 (Table S6). KEGG pathways upregulated in HCM-enriched CM1 and CM4 again exhibited insulin signaling pathways, as well pathways related to mitophagy, focal adhesion/adherens junction, PI3K-Akt signaling, and mTOR signaling (Table S7).

Owing to findings of genes associated with “cardiac stress” in control-enriched clusters, we created module scores incorporating genes identified by Kuppe et al. in “normal”, “pre-stressed”, and “stressed” myocardium derived from adults following myocardial infarctions [20]. Control-enriched clusters CM2 and CM3 had the highest relative expression determined by the “normal” myocardium score, control enriched CM3 and CM5 the highest for the “pre-stressed” myocardium score, and HCM-enriched CM1 and control-enriched CM6 for the “stressed” myocardium score (Figure 2G).

### Co-expression network analysis reveals underlying processes in HCM cardiomyocytes

We then sought to create co-expression networks to identify interconnected genes enriched in HCM by using hdWGCNA. The hdWGCNA pipeline is described in Figure S1A. Twelve modules of co-expressed genes were identified (Figure S1B, Table S8). Correlation of module eigengenes, a metric summarizing gene expression profiles of each module, is exhibited in Figure S1C. Differential module eigengene (DME) analysis results exhibited that module CM-hd9, CM-hd2, and CM-hd7 are upregulated in HCM compared to controls (Figure S1D, Table S9). In CM-hd9, the most upregulated module in HCM, genes with the highest kME include previously implicated *PLCE1* and other genes associated with inflammation, cardiac remodeling, and glucose metabolism like *FKBP5, CPEB4*, and *PDK4* (Figure 1D).[29]–[31] Consistent with our previous analyses, the CM-hd9 hME was highest in HCM-enriched CM1 and CM4 (Figure S1E). Additionally, functional enrichment pathway analysis in CM-hd9 exhibited upregulation of actin cytoskeleton, focal adhesion, and cell substrate junction pathways. CM-hd7 was made up of genes associated with glycolysis, Wnt signaling, and the NF-kappaB (NF- κB) signaling pathways. These pathways were also seen in control dominant CM-hd5 and CM-hd4, implying that pathways related to “cardiac stress” and injury may be seen in structurally normal, donor hearts. Functional enrichment analysis is exhibited in Table S10.

Sensitivity analyses were performed to assess if hdWGCNA findings were consistent with our previous DEG analyses and consistent amongst HCM subjects and cells derived from left or right ventricles. Using overlap analysis and a log threshold of 1 to identify gene markers for CM clusters, we identified significant overlap of CM-hd9 and HCM dominant CM1, with the highest hME expression (Figure S5A-C). CM-hd4, CM-hd11, and CMhd-3 showed variable overlap with “pre-stressed” and “stressed” CM3, CM5, and CM6, which is likely due to genes associated with both scores being housed in those respective modules. We then performed DME on each HCM subject compared to controls and in another analysis only included left ventricle-derived cells, which showed consistent upregulation of CM-hd9 in HCM (Figure S6B, Table S11).

### HCM clusters are enriched in activated fibroblasts

Unbiased re-clustering of fibroblasts identified 4 clusters denoted as FB1-FB4 (Figure 3A). Proportions of cells contributed to each cluster by subject and by total HCM and control are exhibited in Figure 3B. Premutation testing exhibiting a significantly increased proportion of HCM-derived cells in FB3 (Log2FD= 3.27, Adjusted *P-* value<0.001) and increased proportion of control-derived cells in FB1 (Log2FD= 2.06, Adjusted *P-*value<0.001), with a slight predominance of HCM-derived cells in FB2 (Log2FD= 0.45, Adjusted *P-*value<0.001) and FB4 (Log2FD= 0.59, Adjusted *P-* value<0.001). Similar findings were exhibited in the differential abundance analysis, where FB1 was enriched with control neighborhoods and FB3, as well as FB2 and FB4 to a lesser degree, were enriched with HCM neighborhoods (Figure 3C).

**Figure 3:**
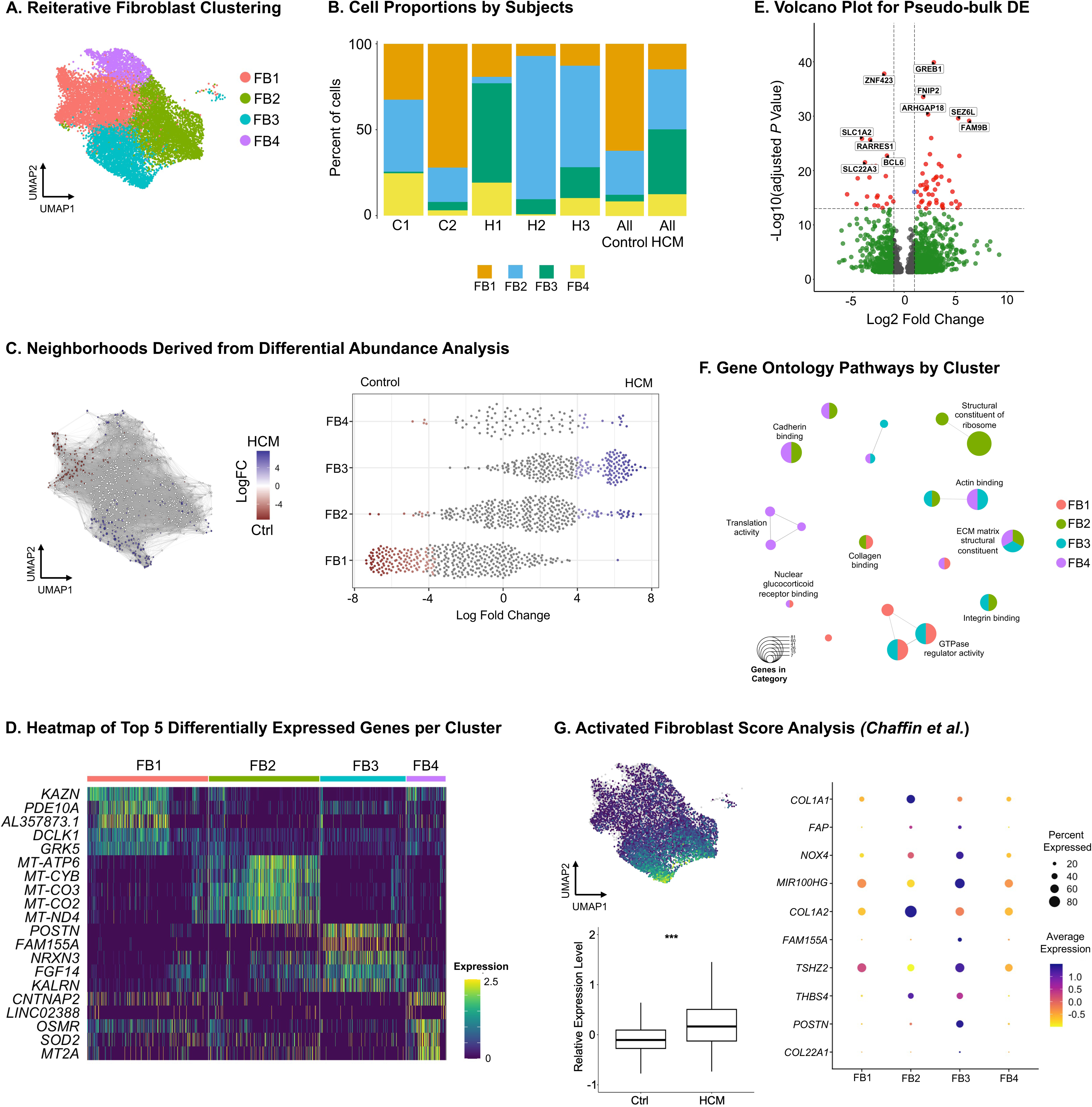
Activated cardiac fibroblast clusters are enriched with cells derived from HCM myocardium. A) UMAP embedding of reiterative clustering of fibroblasts. B) Bar plot depicting proportion of major cell types of interest within control subjects (C1 and C2) and HCM subjects (H1, H2, and H3) and proportions in pooled cohorts (Control vs HCM). C) Left: Embedding derived from Milo k-nearest neighborhood differential abundance testing and layout derived from UMAP embedding. Nodes with false discovery rate <10% depict neighborhoods and are colored by log fold changes for HCM (blue) versus control (red) samples. Right: Beeswarm plot depicting significant log fold changes for HCM (blue) and control (red) neighborhoods derived from differential abundance testing stratified by fibroblast clusters (y-axis). D) Heatmap of top 5 differently expressed genes derived from FindAllMarkers function in FB1, FB2, FB3, and FB4. E) Volcano plot derived from pseudo-bulk RNA differential gene expression. F) Enrichment map stratified by FB cluster depicting the proportional GO pathway analysis. Node size is determined by the number of genes within each GO pathway. G) Top left: Feature plots illustrating expression of activated FB score depicted on original UMAP embedding. Bottom left: Boxplot illustrating relative expression level derived from activated FB score stratified by HCM versus control. The center line in the box represents the median, the top edge of the box shows the upper quartile, the lower edge of the box shows the lower quartile, and the whiskers extend x1.5 of the interquartile range. *P* value <0.001 indicated by ***. Right: Dot plots illustrating average expression (depicted on color scale) and percent of cells expressing each gene (depicted by size of dot), stratified by FB clusters.

In the DEG analysis (Table S12, Figure 3D), HCM-enriched FB3 exhibited DEGs (FC >1) related to fibroblast activation like *POSTN, PALLD, FAM155A,* and *FGF14* [7]. FB2 exhibited marker genes for lipogenic fibroblasts *APOE* and *APOD* [32]. FB4 exhibited pro-inflammatory and pro-fibrotic *OSMR* [33]. This was further validated in pseudo-bulk RNA analysis with significant overlap implicating *POSTN, THSD4, PALLD,* and *FAM155A,* as well as several collagen genes like *COL4A6, COL22A1,* and *COL11A1* (Table S13, Figure 3E). Functional pathway analysis exhibited upregulation of integrin pathways in HCM-enriched FB2 and FB3, extracellular matrix structural constituent and focal adhesion pathways in FB2-FB4, and calmodulin-related pathways in FB3 and FB4 (Table S14-15, Figure 3F). These findings are consistent with fibroblast to myofibroblast activation and their subsequent secretory functions.

To explore the collective expression of genes involved in fibroblast activation, we computed a module score based on the average expression of genes defined in activated fibroblast populations by Chaffin et al.[7] HCM-derived cells exhibited significantly higher module scores and HCM-enriched FB3, consistent with our previous findings, exhibited the highest module score (Figure 3G, Figure S6A). Higher relative expressions for each gene included in the module was shown in HCM compared to controls, as well as higher module scores across all HCM subjects compared to control subjects (Figure S6A). This indicates that our findings are not driven by one HCM subject, but a genetic signature for fibroblast activation across all HCM subjects.

### Activated fibroblast genes are co-expressed and enriched in HCM myocardium

Nine modules of co-expressed genes were identified (Figure S2A; Table S16). Correlation of module eigengenes is exhibited in Figure S2B. Module hMEs are shown overlayed on the original UMAP embedding in Figure S2C. We then applied the UMAP embedding to visualize the co-expression networks as exhibited in Figure S2D. DME analysis results exhibited that modules FB-hd4, FB-hd7, FB CM-hd8 are upregulated in HCM compared to controls (Figure S2E; Table S17). Consistent with our previous DEG analysis, FB-hd4 and FB-hd7 hME was highest in HCM-enriched FB3 (Figure S1F). When focusing on the most upregulated module FB-hd4, fibroblast activating genes *PAALD, POSTN,* and *NOX4* are co-expressed along with genes associated with cell migration, inflammation, and fibrosis like *ASAP1* and *FKBP5* [7],[30],[34]. Similarly, in module FB-hd7, fibroblast activating genes like *THBS4, FAP,* and *LTBP2* are co-expressed with genes associated with fibrosis, cell adhesion, and cell migration like *IGFBP-5*, *FN-1*, and *CD99* [35]–[37]. This implies that fibroblast activation is related to upregulation in cell migration, fibrosis, and inflammation. This is consistent with the functional enrichment pathway analysis showing enrichment of pathways involved in fibrosis in FB-hd7 like extracellular matrix organization pathways and with higher odds ratios compared to control dominant modules (Table S18).

Sensitivity analyses were performed. Overlap analysis exhibited significant overlap between FB-hd4 and FB3 as well as FB-hd7 and FB2, consistent with our previous analyses showing heightened fibroblast activity in those respective clusters (Figure S5A-C). We then performed DME on each HCM subject compared to controls and in another analysis only included left ventricle-derived cells, which showed consistent upregulation of FB-hd4 and FB-hd7 in HCM (Figure S6B, Table S19).

### Genes associated with angiogenesis and cell fate are enriched in HCM endothelial cell and endocardial clusters

Unbiased re-clustering identified five clusters denoted as EC1-EC3, EndoC, and LEC (Figure 4A). Proportions of cells contributed to each cluster by subject and by total HCM and control are exhibited in Figure 4B. With premutation testing of proportions, LECs (Log2FD= 1.29, Adjusted *P-*value=0.002) and EC1 (Log2FD= 0.19, Adjusted *P-* value=0.002) were found to be HCM dominant, while EndoC was control dominant (Log2FD= 1.95, Adjusted *P-*value=0.002). Similarly, differential abundance testing showed that the greatest number of HCM neighborhoods were found in EC1, followed by EC2 and EC3 (Figure 4C).

**Figure 4:**
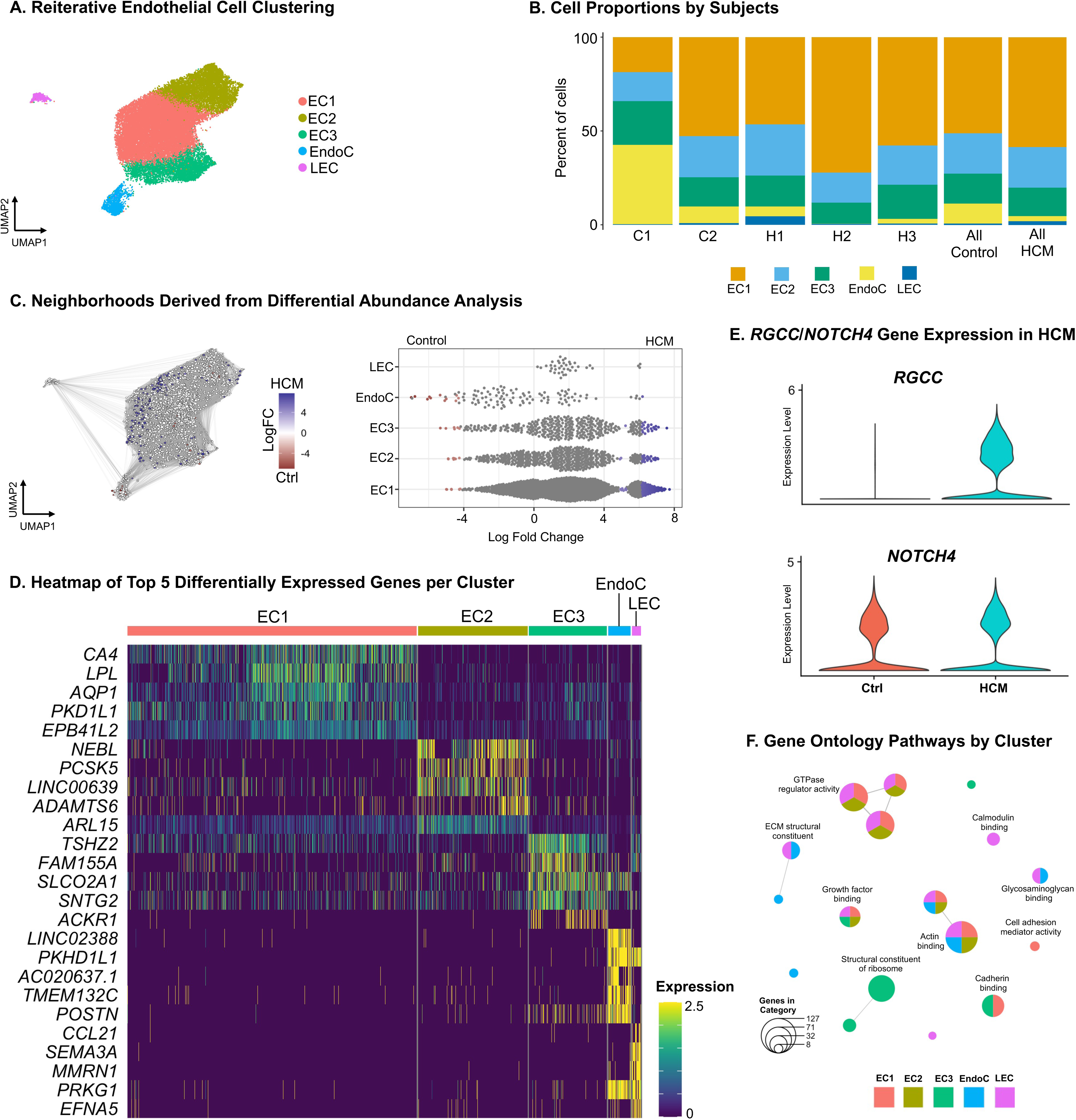
Genes associated with angiogenesis and cell fate are enriched in HCM endothelial cell clusters. A) UMAP embedding of reiterative clustering of endothelial cells. B) Bar plot depicting proportion of major cell types of interest within control subjects (C1 and C2) and HCM subjects (H1, H2, and H3) and proportions in pooled cohorts (Control vs HCM). C) Left: Embedding derived from Milo k-nearest neighborhood differential abundance testing and layout derived from UMAP embedding. Nodes with false discovery rate <10% depict neighborhoods and are colored by log fold changes for HCM (blue) versus control (red) samples. Right: Beeswarm plot depicting significant log fold changes for HCM (blue) and control (red) neighborhoods derived from differential abundance testing stratified by cardiomyocyte clusters (y-axis). D) Heatmap exhibiting differentially expressed genes for each of the EC clusters. Expression levels indicate the average expression for each cluster. Clusters are colored according to Figure 4A. E) Violin plot illustrating relative expression for *RGCC* and *NOTCH4* in HCM versus control cells. F) Enrichment map stratified by EC cluster depicting the proportional GO pathway analysis. Node size is determined by the number of genes within each GO pathway. Node size is determined by the number of genes within each GO pathway.

DEG analysis (Table S20, Figure 4D) was completed. Cluster markers for venous, capillary, and arterial endothelial cells, as well as endocardium and lymphatic endothelial cells were identified. EC1 represented capillary endothelial cells (ie. *CA4* and *RGCC*), EC2 arterial endothelial cells (ie. *GJA5, DLL4,* and *SEMA3G*), EC3 venous endothelial cells (ie. *NR2F2, SELP,* and *ACKR1*), EndoC endocardium (ie. *SPOCK1, NPR3,* and *POSTN*), and LECs lymphatic endothelial cells (ie. *CCL21*).[38] Both EC1 and EC2 were enriched with genes associated with cell fate, angiogenesis, cell junction, and vascular remodeling (*NOTCH4, FLT1, ADAMTS6, FGFR1, RGCC, COL15A1;* Figure 4D/4E) [39]–[41]. This was further validated in pseudo-bulk RNA analysis with overlap implicating genes like *FLT1, FGFR1,* and *RGCC* (Table S21). Functional enrichment pathway analysis exhibited upregulation of GO pathways associated with cell adhesion, cadherin binding, NOTCH binding, and vascular endothelial growth factor receptor binding pathways (Table S22; Figure 4F) and similarly, KEGG pathways associated with adherens junction and NOTCH signaling pathways mainly in clusters EC1-EC3 (Table S23). These findings support the role of endothelial cells in vascular remodeling in HCM.

### Genes driving cell fate and angiogenesis are co-expressed and enriched in HCM

Eight modules of co-expressed genes were identified (Figure S3A; Table S24). Correlation of module eigengenes is exhibited in Figure S3B. Module hMEs are shown overlayed on the original UMAP embedding in Figure S3C. We then applied the UMAP embedding to visualize the co-expression networks as exhibited in Figure S3D. DME analysis results exhibited that module EC-hd2 is upregulated in HCM compared to controls and the hME was highest in HCM-predominant EC1 in our previous DEG analysis (Figure S3E/S3F; Table S25). Hub genes in EC-hd2 again implicate genes associated with angiogenesis like *FLT1*.[39] Consistent with previous results, functional enrichment pathway analysis identified pathways associated with angiogenesis, cell fate, and endothelial cell differentiation (Table S26). Sensitivity analyses were performed similar to prior. Overlap analysis exhibited significant overlap between EC-hd2 and HCM-predominant EC1 and EC2 clusters, of which EC1 had the highest hME expression (Figure S5A-C). We then performed DME on each HCM subject compared to controls and in another analysis only included left ventricle-derived cells, which showed consistent upregulation of EC-hd2 in subjects H1 and H2 and EC-hd2 in HCM compared to LV specific control tissue (Figure S6B, Table S27).

### Tissue resident macrophages are diminished in HCM myocardium

Unbiased re-clustering derived from myeloid cells identified 3 clusters. These were labeled based on cell markers defined by Eraslan et al. as a monocyte derived macrophage cluster enriched with HLA Class II genes and *NAMPT* (MP1), a tissue-resident macrophage cluster enriched with *LYVE1* (MP2), and a monocyte cluster enriched with *VCAN* (Mono) [42]. Proportions of cells contributed to each cluster by subject and by total HCM and control are exhibited in Figure 5B. Premutation testing exhibited heightened HCM proportions in MP1 (Log2FD= 1.21, Adjusted *P-*value<0.001) and heightened control proportions in MP2 (Log2FD= 0.38, Adjusted *P-*value<0.001) and Mono (Log2FD= 2.08, Adjusted *P-*value<0.001). Similar findings were exhibited in the differential abundance analysis, where MP2 and Mono were enriched with control neighborhoods and MP1 more enriched with HCM neighborhoods (Figure 5C).

**Figure 5:**
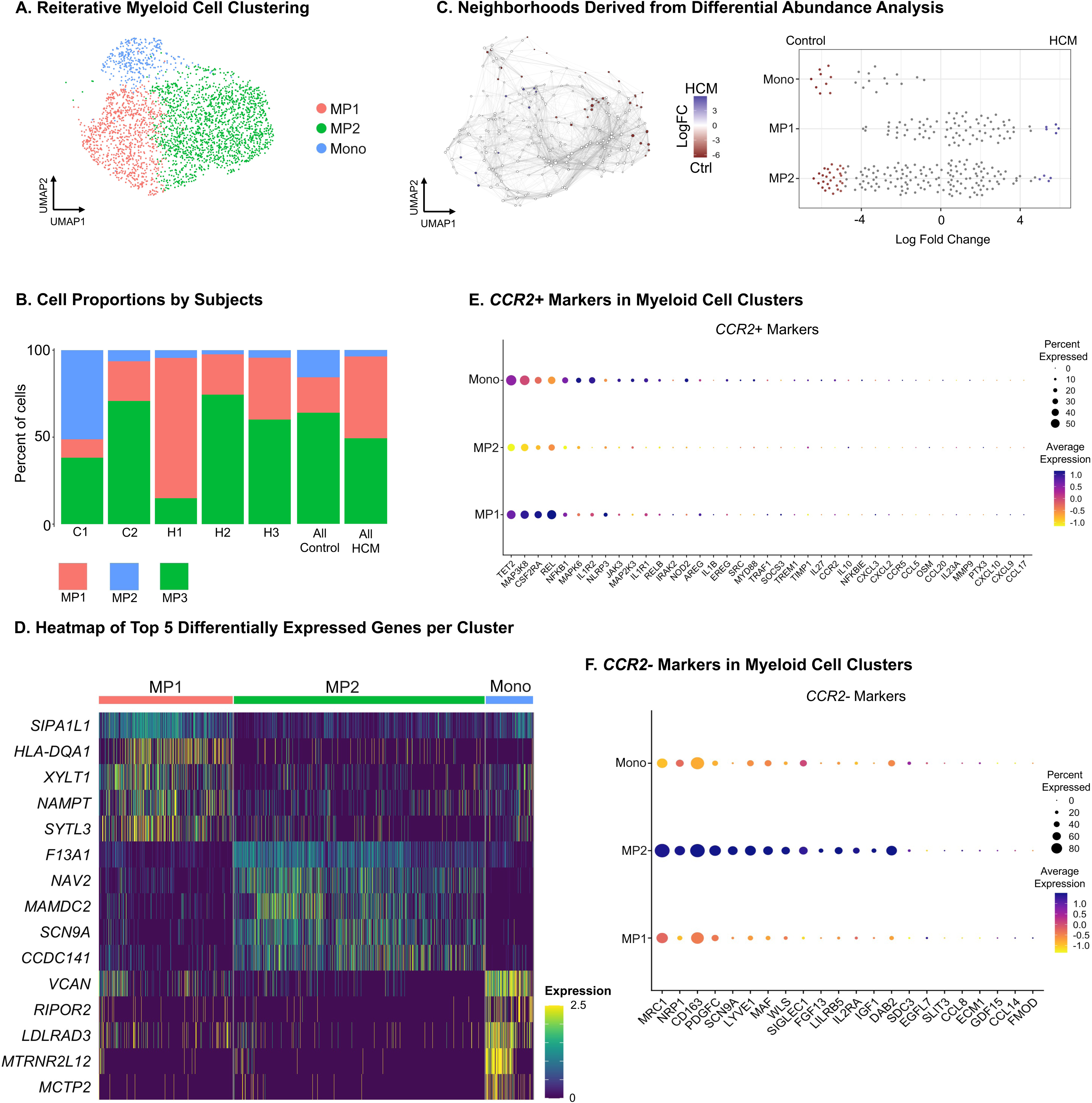
Tissue resident macrophages are predominated by control rather than HCM myocardium. A) UMAP embedding of reiterative clustering of myeloid cells. B) Left: Embedding derived from Milo k-nearest neighborhood differential abundance testing and layout derived from UMAP embedding. Nodes with false discovery rate <10% depict neighborhoods and are colored by log fold changes for HCM (blue) versus control (red) samples. Right: Beeswarm plot depicting significant log fold changes for HCM (blue) and control (red) neighborhoods derived from differential abundance testing stratified by myeloid cell clusters (y-axis). D) Bar plot depicting proportion of major cell types of interest within control subjects (C1 and C2) and HCM subjects (H1, H2, and H3) and proportions in pooled cohorts (Control vs HCM). E) Heatmap exhibiting differentially expressed genes for each of the ML clusters. Expression levels indicate the average expression for each cluster. G) Dot plots illustrating average expression (depicted on color scale) and percent of cells expressing each gene (depicted by size of dot), stratified by ML cells for CCR2+ markers (left) and CCR2- markers (right) derived from Bajpai et al.

DEG analysis (Table S28, Figure 5D) and pseudo-bulk RNA analysis (Table S29) was completed. We also obtained markers of recruited (ie. derived from CCR2+ markers) and resident (derived from CCR2- markers) macrophage subsets identified by Bajpai et al (Figure 5E and 5F) [43]. HCM enriched MP1 differentially expressed (FC>1) *NAMPT,* a gene encoding nicotinamide phosphoribosyltransferase shown to play a role in myocardial adaption to pressure overload, as well as other genes involved in NAD+ nucleosidase activity like *IL18R1* [44]. MP1 also differentially expressed several MHC Class I and II genes (ie. *HLA-DRA, HLA-DRB1, HLA-DQB1, HLA-DPB1*). This is consistent with exhibiting a high relative expression of CCR2+ markers suggesting an inflammatory role for this cluster. This is further substantiated by functional enrichment analysis showing enrichment of GO and KEGG pathways involving antigen processing and presentation, immune activity, and MHC protein-related pathways (Table S30 and S31). MP2 exhibited high relative expression of CCR2- markers and differentially expressed genes like the sodium channel *SCN9A,* a sodium channel modulator *FGF13,* and *F13A1,* encoding the coagulation factor XIII A subunit. This is consistent with findings in previous studies analyzing gene expression in CCR2- macrophages as well as functional enrichment analysis exhibiting pathways associated with scavenger receptor activity and calcium channel activity [43]. When analyzing pseudo-bulk RNA analysis, genes associated with macrophage activation and differentiation were noted, like *ANKRD22* (Table S29) [45].

### HCM myocardium has a predominantly fibroblast driven increase in inferred cellular interactions

We sought to estimate differential outgoing and incoming interactions between HCM and controls. When comparing outgoing and incoming interactions, 86.7% of inferred interactions were in HCM (n=365/421). The differential number and weight of interactions were significantly higher in HCM (Figure 6A-C). When evaluating strength of interactions by cell type, fibroblasts appear to have the highest strength of outgoing interactions while endocardium, endothelial cells, and cardiomyocytes have the highest incoming interaction strengths (Figure 6D). This is consistent with a disrupted tissue microenvironment. Ligand/receptor pairs are shown in Table S32-33. The strongest differential outgoing interactions from fibroblasts involved LAMININ and COLLAGEN pathways and translated to the pathways seen in incoming interactions in cardiomyocytes, endothelial cells, and endocardium (Figure 6E). The ligand and receptors with the highest estimated probabilities include LAMA2, LAMB1, COL6A4, COL4A1, COL4A2, ITGA9, ITGA7, ITGA6, and ITGB1 (Table S32-33). Other outgoing interactions of note include PTPRM pathways, associated with cell growth and differentiation, strongly seen in endocardium and less so in endothelial cells and cardiomyocytes (Figure 6E).

**Figure 6:**
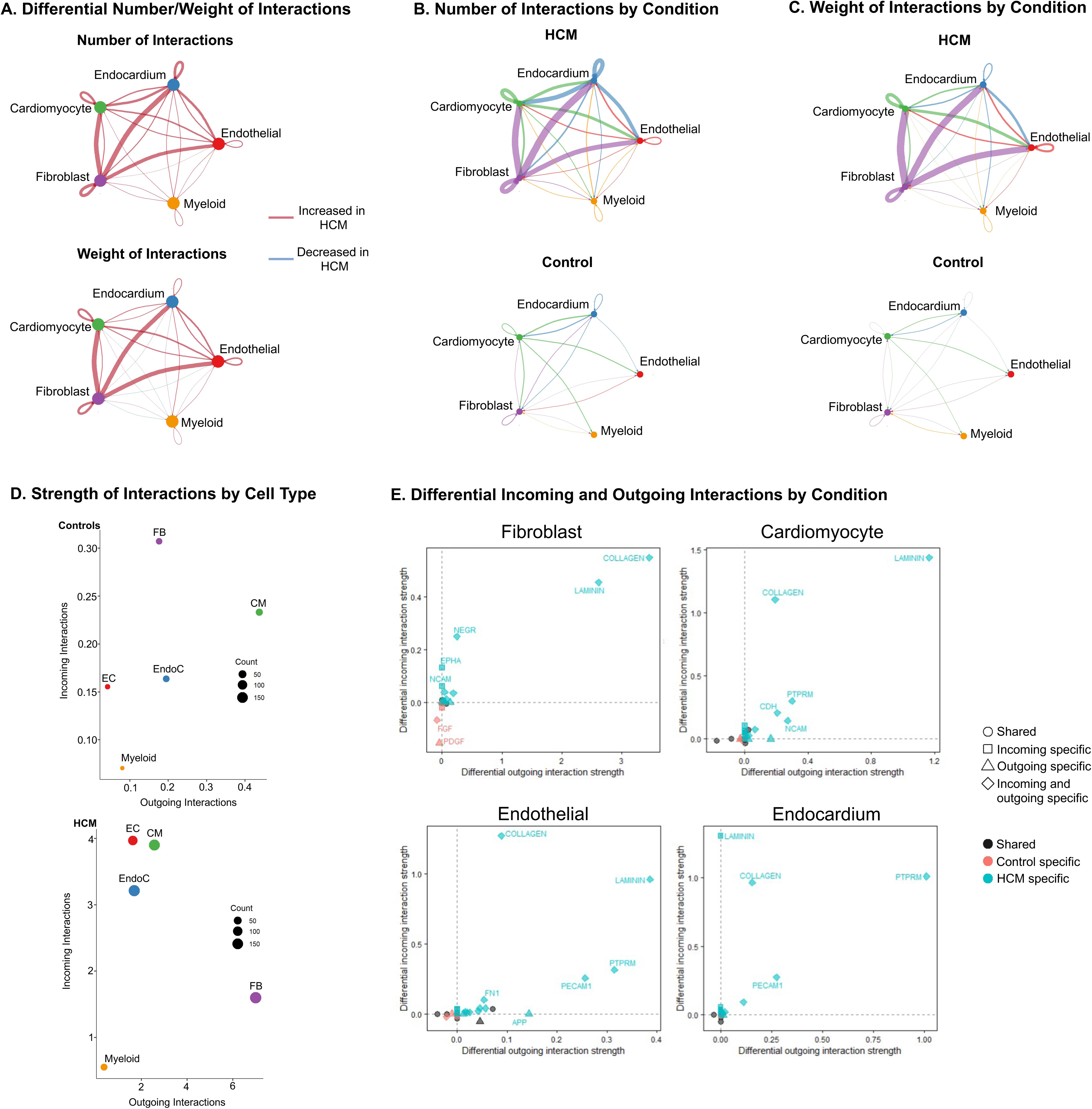
Cell interactions are increased in HCM myocardium and predominantly fibroblast driven. A) Circle plot illustrating the differential number (top) and weight (bottom) of ligand-receptor interactions in HCM versus control cells. Red indicates increased number or weight of interactions in HCM and blue indicates decrease in number or weight of interactions in HCM. B) Circle plots exhibiting the number of ligand-receptor interactions in HCM (top) versus control (bottom) cells. C)) Circle plots exhibiting the weight of ligand-receptor interactions in HCM (top) versus control (bottom) cells. D) Scatter plots for control (top) and HCM (bottom) cells illustrating the strength of incoming (y-axis) and outgoing (x-axis) interactions. The size of each circle is dependent on the number of interactions. E) Scatter plots illustrating differential incoming and outgoing interactions between HCM and control cells in fibroblasts, cardiomyocytes, endothelial cells, and endocardium. Each point represents a signaling pathway. The shapes define if the pathway is shared (circle), incoming specific (square), outgoing specific (triangle), or both incoming and outgoing specific (diamond). The color defines if the pathway is shared (black), control specific (red), or HCM specific (blue).

We then sought to assess cell to cell communication within HCM-derived cells and amongst clusters to identify patterns in diseased tissue. Fibroblast clusters were the largest contributors to both number and strength of outgoing interactions (Figure S4A and S4E) and less so for endothelial cells and endocardium (Figure S4C) and cardiomyocyte clusters (Figure S4D). Ligand/receptor pairs by cluster are shown in Table S34. We then sought to assess common patterns for pathways amongst the clusters. We selected the default 4 patterns to be identified based on Cophenetic and Silhouette values. Outgoing interaction pathways belonging to each pattern are shown in Figure S4B. Cardiomyocyte clusters exhibited pathways in Pattern 1, endothelial clusters EC1-EC3 and EndoC exhibited pathways in Pattern 2, fibroblast clusters in Pattern 3, and LECs in Pattern 4. Specific pathways and their respective contribution in each cluster are shown in Figure S4F and were consistent with pathways expected to be upregulated based on our DEG analyses.

### Pediatric HCM fibroblasts are enriched with ribosomal and extracellular matrix protein-related genes compared to adult HCM

We next integrated data from 16 end-stage, adult HCM subjects described in the Chaffin et al. study [7]. When comparing pediatric HCM to adult HCM, we found 2125 DEGs in pediatric HCM (Table S35). Of these, several DEGs with FC>1 included genes related to ribosomal proteins like *FAU, RPS15, RPL18, and RPL14* and collagen genes like *COL21A1, COL4A1, COL1A1,* and *COL4A4*. In addition, some fibroblast activation genes were upregulated in pediatric HCM like *PALLD, FGF1,* and *AKT3,* but others were upregulated in adult HCM like *FAP, NOX4,* and *THBS4.* Consistent with this, our functional pathway over-representation analysis first on upregulated DEGs in pediatric HCM and then comparing pediatric and adult HCM, exhibited consistent upregulation of GO pathways related to protein synthesis and cellular proliferation (ie. ribosome-related pathways), fibrosis related pathways (ie. extracellular matrix protein-related pathways), and apoptosis/mitophagy pathways (Figure 7A and 7B; Table S36-S38) in pediatric HCM. Owing to heightened fibroblast outgoing interactions noted in our primary analysis, we created module scores based on ligand genes described in CellChat COLLAGEN and LAMININ pathways (https://www.cellchat.org/cellchatdb/). We found consistent upregulation of these modules in pediatric HCM (Figure 7C). This data is consistent with heightened cellular proliferation, protein synthesis, and upregulation of downstream pathways associated with fibrosis.

**Figure 7:**
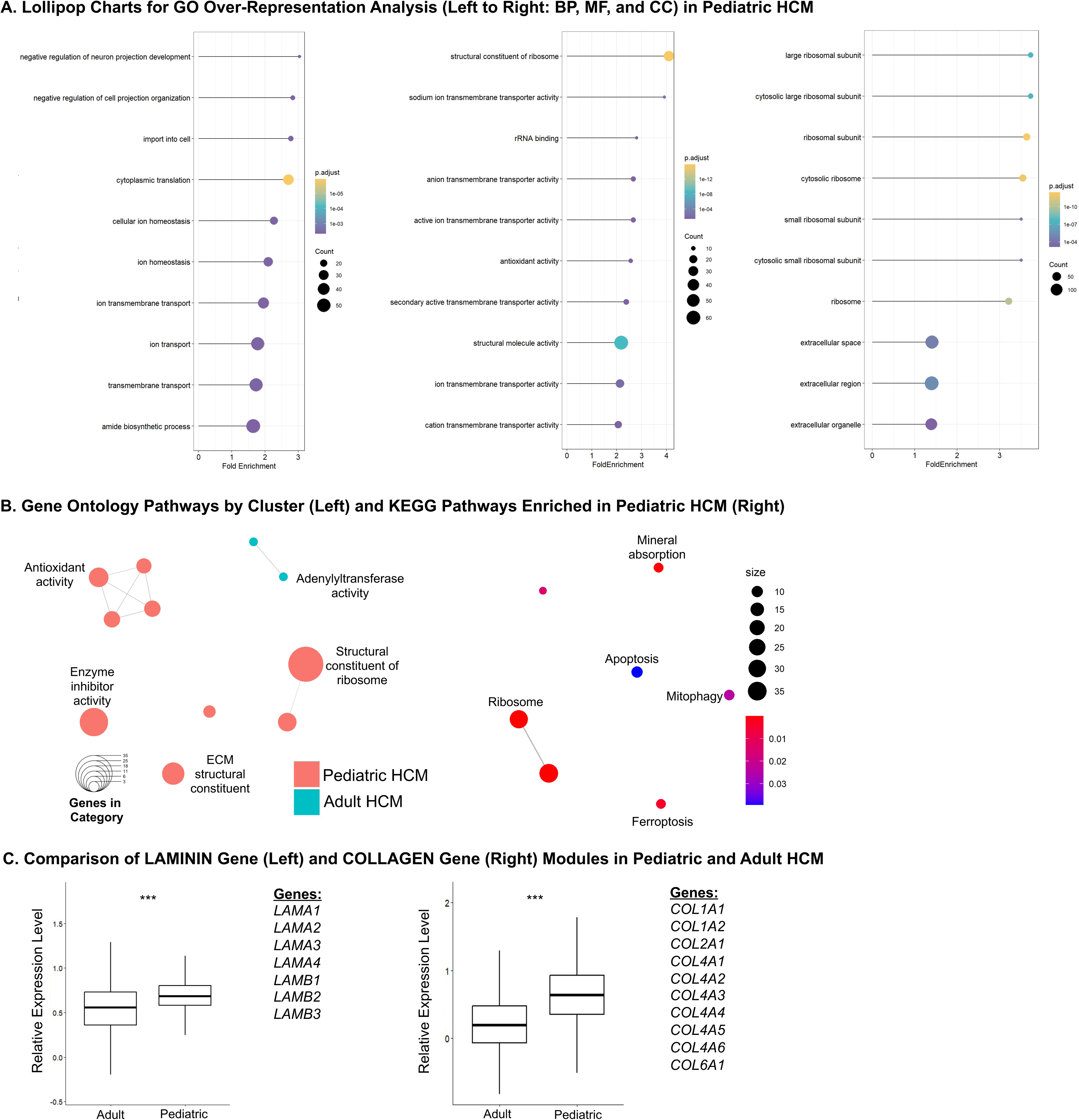
Pediatric HCM are enriched in ECM-related genes compared to adult HCM. A) Lollipop charts for GO over-representation analysis showing top 10 GO pathways (Y-axis) ranked by fold enrichment (X-axis; calculated by mutate(x, FoldEnrichment = parse_ratio(GeneRatio) / parse_ratio(BgRatio))) in biological processes (left), molecular functions (center), and cellular components (right). Colors of circles correspond to adjusted-*P* values and size of circle corresponds to the number of genes implicated. B) Enrichment map stratified by pediatric vs adult HCM depicting the proportional GO pathway analysis (left) and KEGG pathway analysis (right; only pediatric specific pathways met significance). Node size is determined by the number of genes within each pathway. C) Boxplots illustrating relative expression level derived from activated LAMININ module score (left) and COLLAGEN module score (right) stratified by adult versus pediatric HCM. The center line in the box represents the median, the top edge of the box shows the upper quartile, the lower edge of the box shows the lower quartile, and the whiskers extend x1.5 of the interquartile range. *P* value <0.001 indicated by ***.

## Discussion

We identify significant transcriptional changes in ventricular tissue derived from pediatric HCM patients compared to structurally normal hearts in cardiomyocytes, fibroblasts, endothelial/endocardial cells, and myeloid cells. We identify cell states and cellular processes unique to pediatric HCM in major cell types. Interestingly, a marked fibroblast response is seen in end-stage pediatric HCM and a disrupted micro-tissue environment is largely mediated by signaling originating from fibroblasts. Compared to adult HCM, downstream upregulation of fibrotic pathways was seen. These data add to our understanding of end-stage HCM, now uniquely shown in a pediatric population.

We identify genes and pathways upregulated in pediatric HCM cardiomyocytes associated with pathways implicated in hypertrophy, cardiac remodeling, and maladaptive responses to injury. Importantly, we note upregulation of genes associated with “cardiac stress”, like *NPPB* and *ANKRD1* [7],[40]. Interestingly, we note upregulation of other “cardiac stress” genes like *MYH7* and *XIRP2* in some control-enriched clusters. This, as previously shown, may be associated with the organ procurement process [9]. Further, it illustrates that “cardiac stress” can occur outside of structural heart disease. We also identified genes associated with maladaptive and inflammatory responses to stress, cardiac remodeling, and hypertrophy. For instance, *PLCE1* encoding phospholipase C epsilon 1 was identified in both our hdWGCNA and DEG analysis and is known to promote inflammation via NF-κB signaling pathway activation and hypertrophy via scaffolding to mAKAP in cardiac myocytes [26],[46]. Similarly, *FKBP5* was identified which promotes inflammation via NF-κB signaling pathway activation [30]. In addition, other upregulated genes identified included *CPEB4*, a mediator of pathologic cell growth in cardiomyocytes via its interactions with transcription factors, and *PDK4*, of which overexpression has been shown to cause metabolic inflexibility and exacerbate cardiomyopathy in the setting of hypertrophy. [29],[31] We also identified upregulation of pathways in HCM-enriched clusters that are key regulators of hypertrophy and can cross-interact to promote hypertrophy, like insulin signaling pathways, PI3K-Akt signaling, and mTOR signaling [47].

Fibroblasts are the main mediators of fibrosis in heart disease and cardiac fibroblasts have the capabilities to expand functionally in response to injury [48]. Although their functions can be reparative in the acute setting, chronic fibroblast activation and subsequent fibrosis can lead to diastolic and systolic dysfunction, as well as have pro-arrhythmic consequences. Clinically, fibrotic remodeling seen in cardiac disease, particularly in heart failure with preserved ejection fraction, is associated with poorer outcomes [49]. Activated fibroblasts, or myofibroblasts, are usually seen in the early proliferative phase shortly after myocardial injury and this population decreases as the scar matures [48]. They possess several functional roles including neovessel and extracellular matrix formation, as well as contractile activity [48]. We note a large population of activated fibroblasts in end-stage pediatric HCM. Fibroblast activation is noted in adult HCM studies as well and were almost exclusively found in pathologic samples [7]. This has also been exhibited in adult DCM, with robust evidence of activation gene signatures [12]. When comparing our pediatric data to adult HCM fibroblast data, we found significant upregulation of pathways associated with cellular proliferation and protein synthesis. Further, although fibroblast activation genes were not consistently upregulated in pediatric HCM, we found downstream pathways associated with fibrosis (ie. ECM-related pathways) were upregulated in pediatric HCM compared to adults. This may explain why these patients presented with restrictive physiology early on and met indication for transplant prior to their adult counterparts.

Other notable findings in non-cardiomyocyte cell types are a predominance of endothelial cell heterogeneity and enrichment in HCM, as well as a depletion of tissue-resident macrophages. In HCM, myocardial ischemia, especially in the subendocardium, can be seen in the absence of coronary artery disease and is likely secondary to a cardiac mass/perfusion mismatch [50]. As such, responses seen in endothelial cells promoting angiogenesis is likely a response to chronically hypoxic myocardium and an attempt to maintain contractile function [51]. Although the exact mechanisms are not fully understood, angiogenesis-induced hypertrophy can still occur without hypertrophic stimulation [51]. In addition, pathways related to angiogenesis like the NOTCH and vascular endothelial growth factor receptor binding pathways, were found to be upregulated [51],[52]. In regards to myeloid cells, the resident marker enriched cluster was deplete of HCM neighborhoods and the recruited marker rich cluster had slightly higher amounts of HCM neighborhoods. Resident macrophages, enriched with CCR2- markers, are protective and compared to more pro-inflammatory CCR2+ macrophages, do not secrete inflammatory mediators and may inhibit leukocyte recruitment [43]. This may suggest a shift to a more pro-inflammatory myeloid state in pediatric HCM.

Using ligand to receptor analyses, we identify a clear upregulation of cell-to-cell interactions in HCM compared to controls. This is consistent with a disrupted microtissue environment and aberrant intercellular communication seen in disease [9]. On further analysis, the majority of outgoing interactions are noted to be mediated by fibroblasts and were mostly related to collagen and laminin pathways, with cardiomyocytes, endocardium, and endothelial cells being the primary receivers of the interactions. The primary receptors identified were of the integrin family, a group of heterodimeric transmembrane cell adhesion molecules, that likely play a role in extracellular matrix protein deposition in the diseased heart [53]. As pathologic remodeling continues, mechanical stiffness in of itself can promote myofibroblast differentiation and continued extracellular matrix protein production [53],[54]. Our overall data suggests a prominent fibroblast-mediated response leading to extracellular matrix protein deposition.

Our study is limited by the sample size and lack of functional validation. Pediatric samples and disease are relatively rare compared to their adult counterparts and as such, this will be an inherent limitation to most pediatric-focused studies. In addition, biases that cannot be controlled for in our patients like race, ethnicity, surgical sampling, medication, and epi-genetic modifications may have influence on our results. And finally, patients used in this study are those with end-stage disease requiring HT and as such, may not represent cellular processes seen in non-end-stage disease. Regardless, this is the first snRNA-seq study seeking to illustrate underling genetic signatures in pediatric HCM.

In conclusion, our data exhibits prominent underlying cellular processes in pediatric HCM. Fibroblast-mediated cellular processes appear to be the most prominent amongst the major cardiac cell types analyzed and when compared to adult counterparts, pediatric patients transplanted appear to have a heightened amount of downstream processes associated with fibrosis.

## Figure Legends

**Figure S1:**
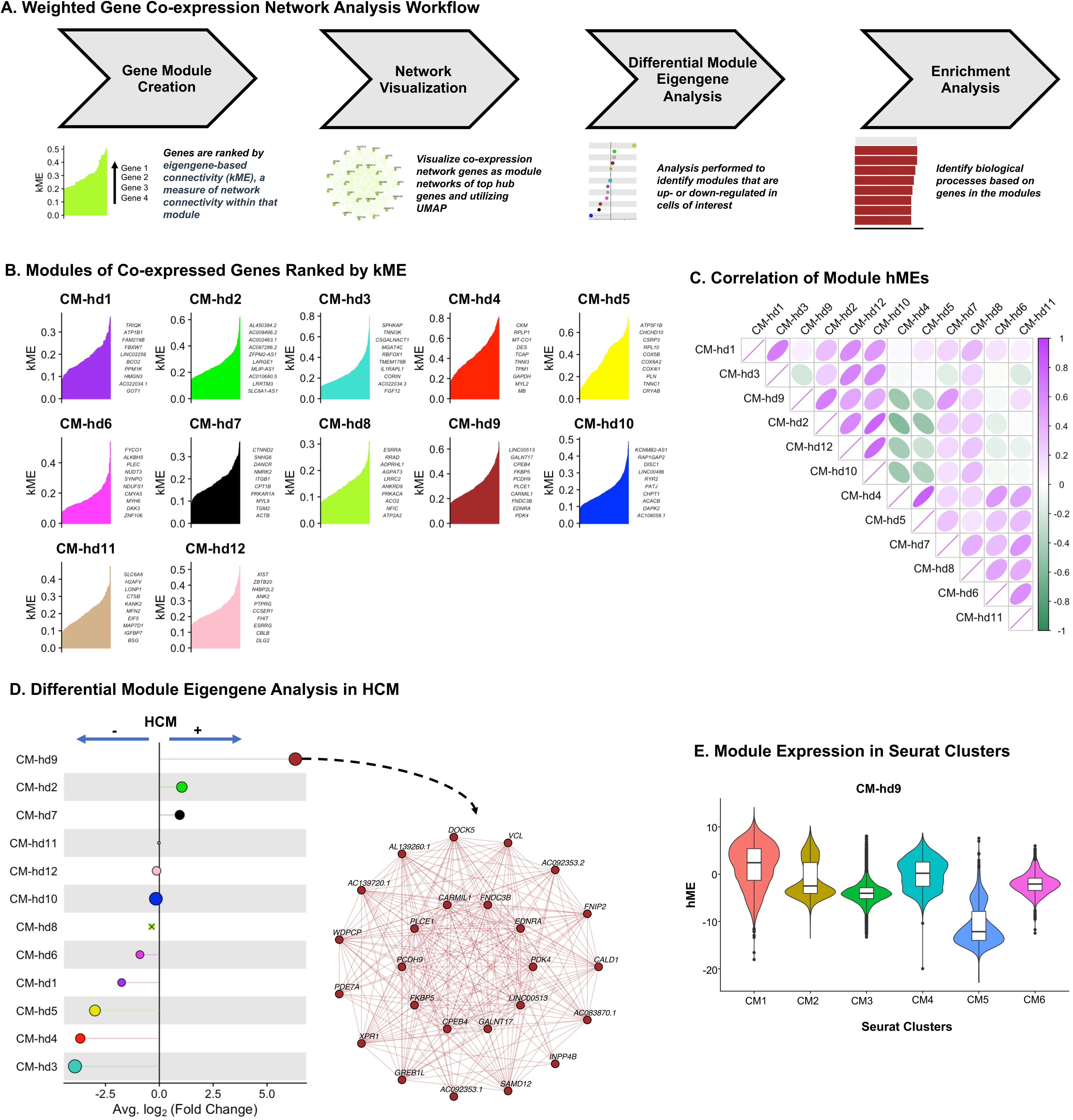
Co-expression network analysis reveals underlying processes in HCM cardiomyocytes. A) High definition weighted gene co-expression network analysis workflow. B) Plots exhibiting genes in each module ranked by e*igengene-based connectivity* (kME). C) Correlogram of module eigengenes, a metric summarizing gene expression profiles of co-expression modules, with batch correction completed with Harmony (hMEs). Positive correlation is indicated by a positive numeric and a purple color and negative correlation is indicated by a negative numeric and a green color. Non-significant correlations are indicated by an “X”. D) Left: Lollipop plot of differential module eigengene (DME) analysis results, illustrating average log2 fold change for each of the modules within HCM. The size of each dot corresponds to the number of genes in the module. Non-significant correlations are indicated by an “X”. Right: Network module plot for upregulated CM-hd9 exhibiting the top 25 genes with the highest kME. E) Violin plot for upregulated module CM-hd9 hME stratified by Seurat FB clusters, with box plots overlayed. The center line in the box represents the median, the top edge of the box shows the upper quartile, the lower edge of the box shows the lower quartile, and the whiskers extend x1.5 of the interquartile range.

**Figure S2:**
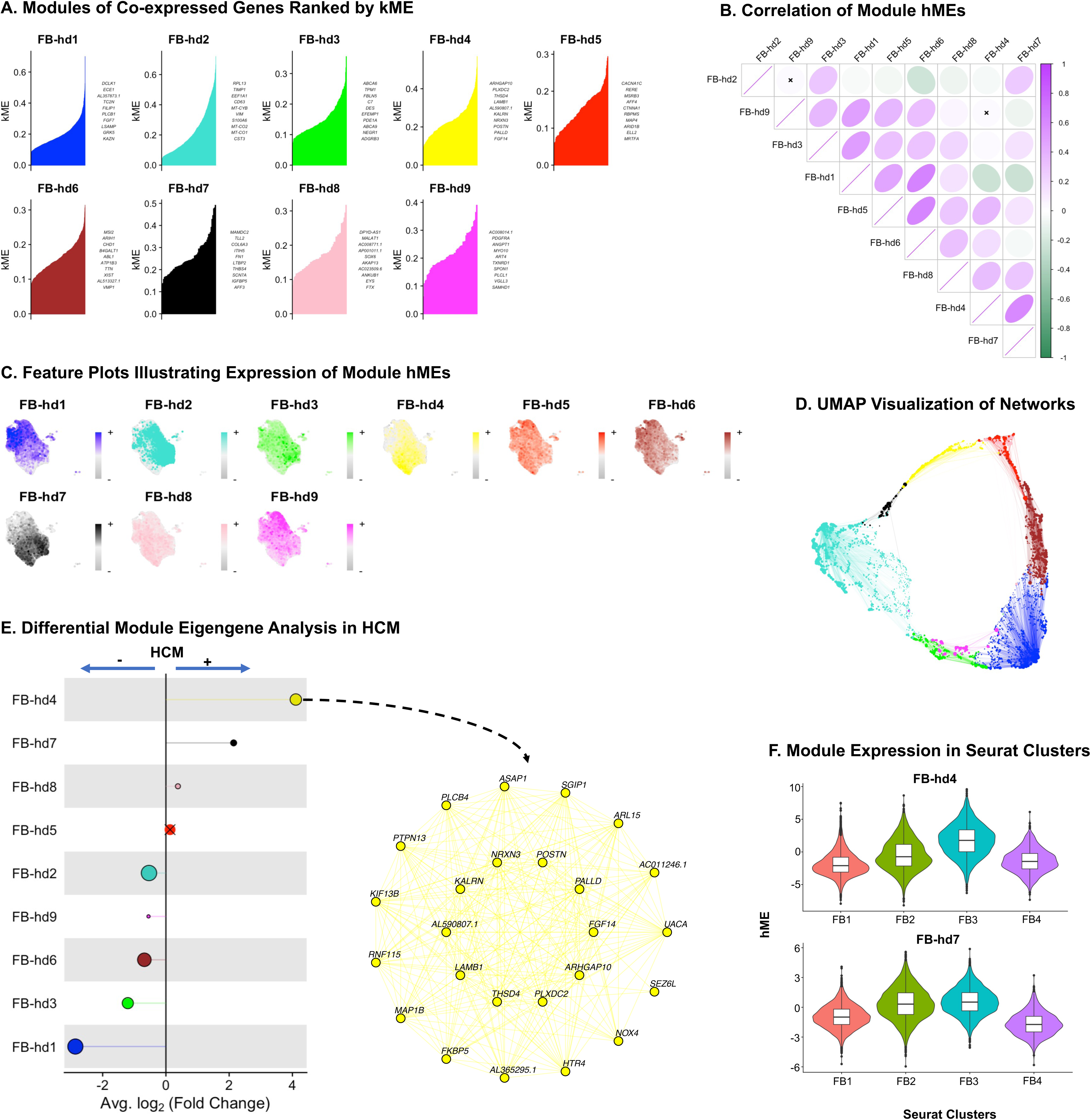
Activated fibroblast genes are co-expressed and enriched in HCM myocardium. A) Plots exhibiting genes in each module ranked by e*igengene-based connectivity* (kME). B) Correlogram of module eigengenes, a metric summarizing gene expression profiles of co-expression modules, with batch correction completed with Harmony (hMEs). Positive correlation is indicated by a positive numeric and a purple color and negative correlation is indicated by a negative numeric and a green color. Non-significant correlations are indicated by an “X”. C) Feature plot created within hdWGCNA illustrating expression of module hMEs colored by each module’s assigned color, and depicted on the original UMAP embedding. D) UMAP embedding applied on the hdWGCNA network to visualize the co-expression network, containing the top 10 hub genes stratified by kME for each module, and downsampled to keep only 20% of edges in the network for visual clarity. Points represent each gene and the size of each dot is scaled based on a gene’s kME for its respective module. E) Left: Lollipop plot of differential module eigengene (DME) analysis results, illustrating average log2 fold change for each of the modules within HCM. The size of each dot corresponds to the number of genes in the module. Non-significant correlations are indicated by an “X”. Right: Network module plot for upregulated FB-hd4 exhibiting the top 25 genes with the highest kME. E) Violin plot for upregulated module FB-hd4 and FB-hd7 hMEs stratified by Seurat FB clusters, with box plots overlayed. The center line in the box represents the median, the top edge of the box shows the upper quartile, the lower edge of the box shows the lower quartile, and the whiskers extend x1.5 of the interquartile range.

**Figure S3:**
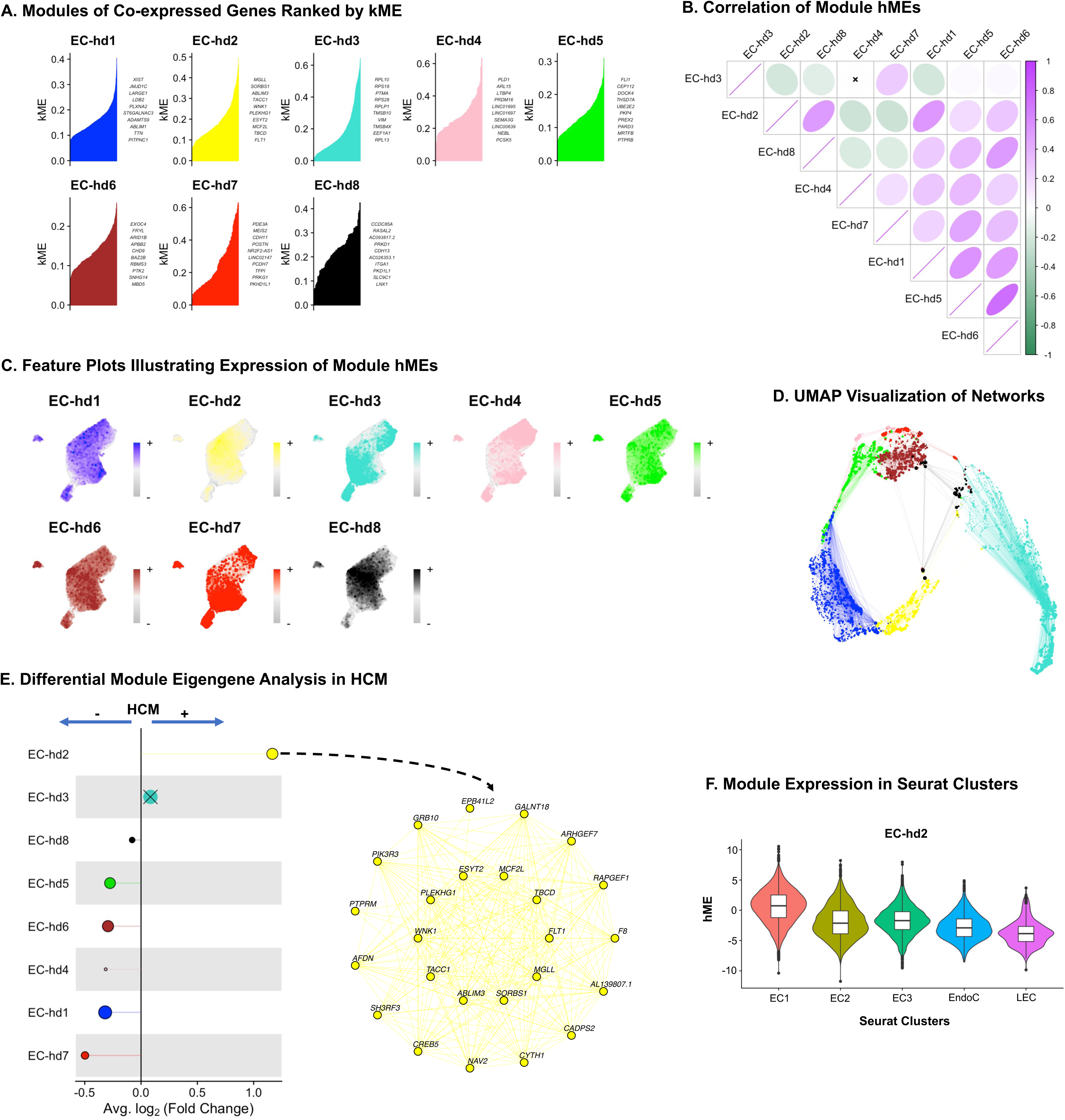
Genes driving cell fate and angiogenesis are co-expressed and enriched in HCM. A) Plots exhibiting genes in each module ranked by e*igengene-based connectivity* (kME). B) Correlogram of module eigengenes, a metric summarizing gene expression profiles of co-expression modules, with batch correction completed with Harmony (hMEs). Positive correlation is indicated by a positive numeric and a purple color and negative correlation is indicated by a negative numeric and a green color. Non-significant correlations are indicated by an “X”. C) Feature plot created within hdWGCNA illustrating expression of module hMEs colored by each module’s assigned color, and depicted on the original UMAP embedding. D) UMAP embedding applied on the hdWGCNA network to visualize the co-expression network, containing the top 10 hub genes stratified by kME for each module, and downsampled to keep only 20% of edges in the network for visual clarity. Points represent each gene and the size of each dot is scaled based on a gene’s kME for its respective module. E) Left: Lollipop plot of differential module eigengene (DME) analysis results, illustrating average log2 fold change for each of the modules within HCM. The size of each dot corresponds to the number of genes in the module. Non-significant correlations are indicated by an “X”. Right: Network module plot for upregulated EC-hdd exhibiting the top 25 genes with the highest kME. E) Violin plot for upregulated module EC-hd2 hME stratified by Seurat FB clusters, with box plots overlayed. The center line in the box represents the median, the top edge of the box shows the upper quartile, the lower edge of the box shows the lower quartile, and the whiskers extend x1.5 of the interquartile range.

**Figure S4:**
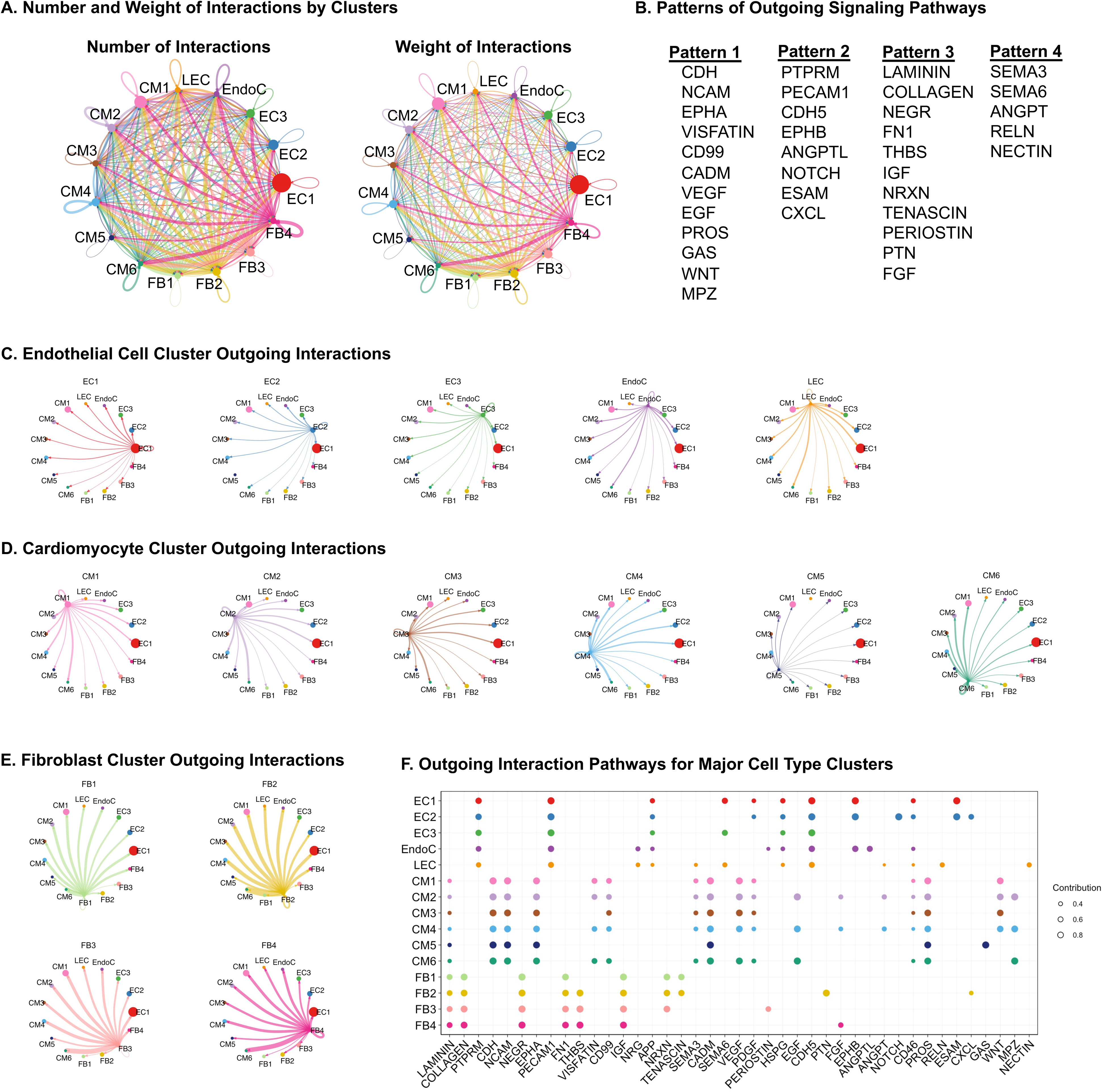
Outgoing interactions are driven by fibroblast cluster 3 and 4. A) Circle plot illustrating the number (left) and weight (right) of ligand-receptor interactions in HCM cells within each fibroblast, cardiomyocyte, endocardium, and endothelial cell cluster. B) Outgoing patterns showing how sender cells drive communication with certain signaling pathways. Four patterns were identified with their respective signaling pathways. Fibroblasts were associated with Pattern 3, cardiomyocytes with Pattern 1, endothelial cells with Pattern 2, and endocardium with Pattern 4. C)-E) Circle plots for each endothelial, endocardial, cardiomyocyte, and fibroblast cluster illustrating the relative weights of outgoing interactions. F) Dot plots illustrating outgoing communication patterns from each of the individual clusters (y-axis) and the respective signaling pathways (x-axis). The size of each dot is representative of the relative contribution.

**Figure S5:**
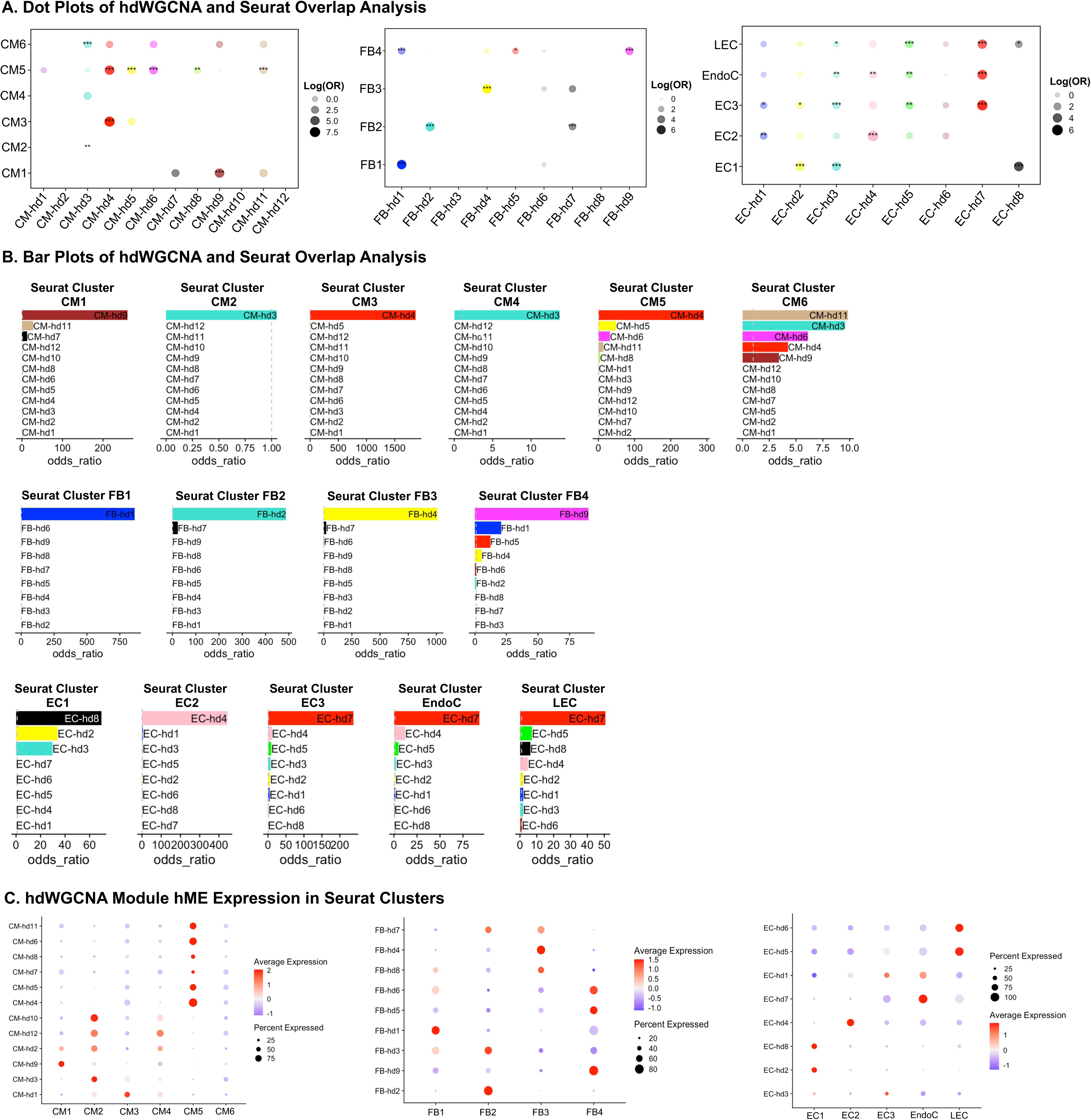
Modules of interest derived from hdWGCNA are enriched in Seurat clusters of interest. A) Dot plots illustrating the results of the cardiomyocyte (left), fibroblast (center), and endothelial cell/endocardium (right) hdWGCNA and Seurat overlap analyses. The presence of each dot represents overlap of our Seurat marker genes and the respective hdWGCNA module. The size of each dot represents the overlap analysis statistic. The false discovery rate (FDR) is represented as *** for FDR of 0-0.001, ** for FDR of 0.001-0.01, and * for FDR of 0.01-0.05. B) Bar plots illustrating the results of the cardiomyocyte (top), fibroblast (middle), and endothelial cell/endocardium (bottom) hdWGCNA and Seurat overlap analysis. hdWGCNA modules are ranked by the overlap statistic. A grey line illustrates an odds ratio of 1. C) Bar plot illustrating the results of the cardiomyocyte hdWGCNA and Seurat overlap analysis. hdWGCNA modules are ranked by the overlap statistic. A grey line illustrates a FDR <0.05. D) Dot plots exhibiting harmonized module eigengenes (hMEs) for each respective hdWGCNA module within the Seurat clusters in cardiomyocyte (left), fibroblast (center), and endothelial cell/endocardium (right). The average expression is depicted on a color scale and percent of cells expressing each gene is depicted by size of the dot.

**Figure S6:**
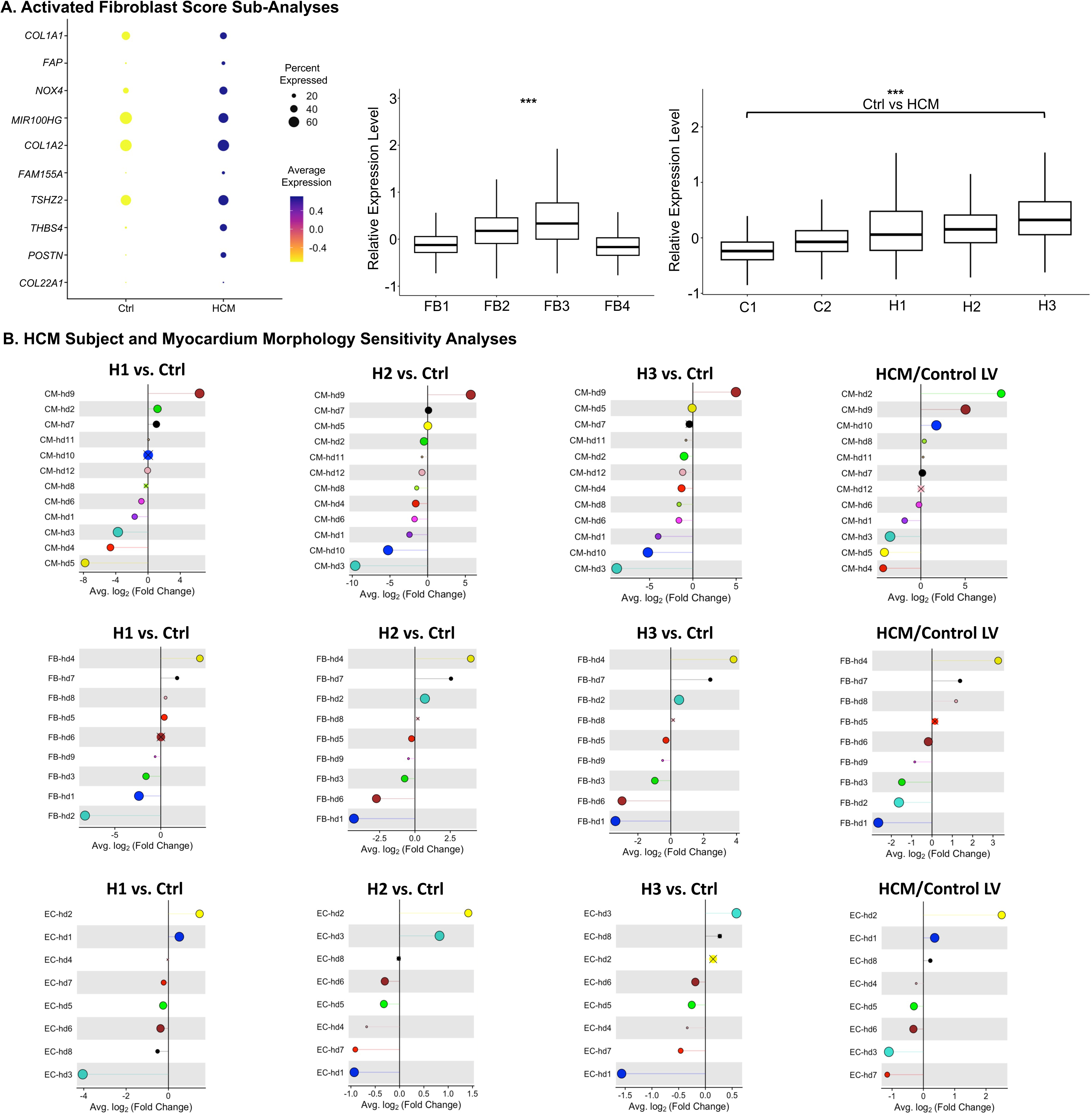
Additional sensitivity analyses exhibits consistency amongst HCM subjects. A) Left: Dot plot illustrating average expression (depicted on color scale) and percent of cells expressing each gene (depicted by size of dot), stratified by HCM versus Control. Center: Boxplots illustrating relative expression level derived from activated FB score stratified by stratified by FB clusters (center). Right: Boxplots illustrating relative expression level derived from activated FB score stratified by individual subjects. The center line in the box represents the median, the top edge of the box shows the upper quartile, the lower edge of the box shows the lower quartile, and the whiskers extend x1.5 of the interquartile range. *P* value <0.001 indicated by ***. B) Lollipop plots illustrating the DME analysis results in the cardiomyocyte, fibroblast, and endothelial hdWGCNA analysis when each HCM subject is compared against control subjects independently and when only LV to LV tissue is analyzed.

## References

1. Arola A, Jokinen E, Ruuskanen O, Saraste M, Pesonen E, Kuusela AL, Tikanoja T, Paavilainen T, Simell O. Epidemiology of Idiopathic Cardiomyopathies in Children and Adolescents: A Nationwide Study in Finland. Am J Epidemiol. 1997;146(5):385–393. doi:10.1093/oxfordjournals.aje.a009291

2. Norrish G, Field E, Mcleod K, Ilina M, Stuart G, Bhole V, Uzun O, Brown E, Daubeney PEF, Lota A, Linter K, Mathur S, Bharucha T, Kok KL, Adwani S, Jones CB, Reinhardt Z, Kaski JP. Clinical presentation and survival of childhood hypertrophic cardiomyopathy: a retrospective study in United Kingdom. Eur Heart J. 2019;40(12):986–993. doi:10.1093/eurheartj/ehy798

3. Ho CY, Day SM, Ashley EA, Michels M, Pereira AC, Jacoby D, Cirino AL, Fox JC, Lakdawala NK, Ware JS, Caleshu CA, Helms AS, Colan SD, Girolami F, Cecchi F, Seidman CE, Sajeev G, Signorovitch J, Green EM, Olivotto I. Genotype and Lifetime Burden of Disease in Hypertrophic Cardiomyopathy: Insights from the Sarcomeric Human Cardiomyopathy Registry (SHaRe). Circulation. 2018;138(14):1387–1398. doi:10.1161/CIRCULATIONAHA.117.033200

4. Gajarski R, Naftel DC, Pahl E, Alejos J, Pearce FB, Kirklin JK, Zamberlan M, Dipchand AI. Outcomes of pediatric patients with hypertrophic cardiomyopathy listed for transplant. J Hear lung Transplant Off Publ Int Soc Hear Transplant. 2009;28(12):1329–1334. doi:10.1016/j.healun.2009.05.028

5. Marian AJ, Braunwald E. Hypertrophic Cardiomyopathy. Circ Res. 2017;121(7):749-770. doi:10.1161/CIRCRESAHA.117.311059

6. Tadros HJ, Life CS, Garcia G, Pirozzi E, Jones EG, Datta S, Parvatiyar MS, Chase PB, Allen HD, Kim JJ, Pinto JR, Landstrom AP. Meta-analysis of cardiomyopathy-associated variants in troponin genes identifies loci and intragenic hot spots that are associated with worse clinical outcomes. J Mol Cell Cardiol. 2020;142:118–125. doi:10.1016/j.yjmcc.2020.04.005

7. Chaffin M, Papangeli I, Simonson B, Akkad AD, Hill MC, Arduini A, Fleming SJ, Melanson M, Hayat S, Kost-Alimova M, Atwa O, Ye J, Bedi KCJ, Nahrendorf M, Kaushik VK, Stegmann CM, Margulies KB, Tucker NR, Ellinor PT. Single-nucleus profiling of human dilated and hypertrophic cardiomyopathy. Nature. 2022;608(7921):174-180. doi:10.1038/s41586-022-04817-8

8. Liu X, Yin K, Chen L, Chen W, Li W, Zhang T, Sun Y, Yuan M, Wang H, Song Y, Wang S, Hu S, Zhou Z. Lineage-specific regulatory changes in hypertrophic cardiomyopathy unraveled by single-nucleus RNA-seq and spatial transcriptomics. Cell Discov 2023 91. 2023;9(1):1-25. doi:10.1038/s41421-022-00490-3

9. Hill MC, Kadow ZA, Long H, Morikawa Y, Martin TJ, Birks EJ, Campbell KS, Nerbonne J, Lavine K, Wadhwa L, Wang J, Turaga D, Adachi I, Martin JF. Integrated multi-omic characterization of congenital heart disease. Nature. 2022;608(7921):181-191. doi:10.1038/s41586-022-04989-3

10. Mehdiabadi NR, Boon Sim C, Phipson B, Kalathur RKR, Sun Y, Vivien CJ, ter Huurne M, Piers AT, Hudson JE, Oshlack A, Weintraub RG, Konstantinov IE, Palpant NJ, Elliott DA, Porrello ER. Defining the Fetal Gene Program at Single-Cell Resolution in Pediatric Dilated Cardiomyopathy. Circulation. 2022;146(14):1105–1108. doi:10.1161/CIRCULATIONAHA.121.057763

11. Hill MC, Martin JF. Epigenetic Assays in Purified Cardiomyocyte Nuclei. Methods Mol Biol. 2021;2158:307–321. doi:10.1007/978-1-0716-0668-1_23

12. Koenig AL, Shchukina I, Amrute J, Andhey PS, Zaitsev K, Lai L, Bajpai G, Bredemeyer A, Smith G, Jones C, Terrebonne E, Rentschler SL, Artyomov MN, Lavine KJ. Single-cell transcriptomics reveals cell-type-specific diversification in human heart failure. Nat Cardiovasc Res. 2022;1(3):263–280. doi:10.1038/s44161-022-00028-6

13. Wolock SL, Lopez R, Klein AM. Scrublet: Computational Identification of Cell Doublets in Single-Cell Transcriptomic Data. Cell Syst. 2019;8(4):281–291.e9. doi:10.1016/j.cels.2018.11.005

14. Xu C, Lopez R, Mehlman E, Regier J, Jordan MI, Yosef N. Probabilistic harmonization and annotation of single-cell transcriptomics data with deep generative models. Mol Syst Biol. 2021;17(1):e9620. doi:10.15252/msb.20209620

15. Love MI, Huber W, Anders S. Moderated estimation of fold change and dispersion for RNA-seq data with DESeq2. Genome Biol. 2014;15(12):550. doi:10.1186/s13059-014-0550-8

16. Dann E, Henderson NC, Teichmann SA, Morgan MD, Marioni JC. Differential abundance testing on single-cell data using k-nearest neighbor graphs. Nat Biotechnol. 2022;40(2):245–253. doi:10.1038/s41587-021-01033-z

17. Langfelder P, Horvath S. WGCNA: an R package for weighted correlation network analysis. BMC Bioinformatics. 2008;9:559. doi:10.1186/1471-2105-9-559

18. Morabito S, Reese F, Rahimzadeh N, Miyoshi E, Swarup V. hdWGCNA identifies co-expression networks in high-dimensional transcriptomics data. Cell reports methods. 2023;3(6):100498. doi:10.1016/j.crmeth.2023.100498

19. Morabito S, Miyoshi E, Michael N, Shahin S, Martini AC, Head E, Silva J, Leavy K, Perez-Rosendahl M, Swarup V. Single-nucleus chromatin accessibility and transcriptomic characterization of Alzheimer’s disease. Nat Genet. 2021;53(8):1143–1155. doi:10.1038/s41588-021-00894-z

20. Kuppe C, Ramirez Flores RO, Li Z, Hayat S, Levinson RT, Liao X, Hannani MT, Tanevski J, Wünnemann F, Nagai JS, Halder M, Schumacher D, Menzel S, Schäfer G, Hoeft K, Cheng M, Ziegler S, Zhang X, Peisker F, Kaesler N, Saritas T, Xu Y, Kassner A, Gummert J, Morshuis M, Amrute J, Veltrop RJA, Boor P, Klingel K, Van Laake LW, Vink A, Hoogenboezem RM, Bindels EMJ, Schurgers L, Sattler S, Schapiro D, Schneider RK, Lavine K, Milting H, Costa IG, Saez-Rodriguez J, Kramann R. Spatial multi-omic map of human myocardial infarction. Nature. 2022;608(7924):766-777. doi:10.1038/s41586-022-05060-x

21. Yu G, Wang LG, Han Y, He QY. clusterProfiler: an R package for comparing biological themes among gene clusters. OMICS. 2012;16(5):284–287. doi:10.1089/omi.2011.0118

22. Jin S, Guerrero-Juarez CF, Zhang L, Chang I, Ramos R, Kuan CH, Myung P, Plikus M V, Nie Q. Inference and analysis of cell-cell communication using CellChat. Nat Commun. 2021;12(1):1088. doi:10.1038/s41467-021-21246-9

23. Bunis DG, Andrews J, Fragiadakis GK, Burt TD, Sirota M. dittoSeq: universal user-friendly single-cell and bulk RNA sequencing visualization toolkit. Bioinformatics. 2021;36(22-23):5535–5536. doi:10.1093/bioinformatics/btaa1011

24. Miller SA, Policastro RA, Sriramkumar S, Lai T, Huntington TD, Ladaika CA, Kim D, Hao C, Zentner GE, O’Hagan HM. LSD1 and Aberrant DNA Methylation Mediate Persistence of Enteroendocrine Progenitors That Support BRAF-Mutant Colorectal Cancer. Cancer Res. 2021;81(14):3791–3805. doi:10.1158/0008-5472.CAN-20-3562

25. Diguet N, Trammell SAJ, Tannous C, Deloux R, Piquereau J, Mougenot N, Gouge A, Gressette M, Manoury B, Blanc J, Breton M, Decaux JF, Lavery GG, Baczkó I, Zoll J, Garnier A, Li Z, Brenner C, Mericskay M. Nicotinamide riboside preserves cardiac function in a mouse model of dilated cardiomyopathy. Circulation. 2018;137(21):2256–2273. doi:10.1161/CIRCULATIONAHA.116.026099/-/DC1

26. Li WH, Li Y, Chu Y, Wu WM, Yu QH, Zhu XB, Wang Q. PLCE1 promotes myocardial ischemia– reperfusion injury in H/R H9c2 cells and I/R rats by promoting inflammation. Biosci Rep. 2019;39(7):20181613. doi:10.1042/BSR20181613

27. Abdellatif M, Trummer-Herbst V, Heberle AM, Humnig A, Pendl T, Durand S, Cerrato G, Hofer SJ, Islam M, Voglhuber J, Ramos Pittol JM, Kepp O, Hoefler G, Schmidt A, Rainer PP, Scherr D, Von Lewinski D, Bisping E, McMullen JR, Diwan A, Eisenberg T, Madeo F, Thedieck K, Kroemer G, Sedej S. Fine-Tuning Cardiac Insulin-Like Growth Factor 1 Receptor Signaling to Promote Health and Longevity. Circulation. 2022;145(25):1853–1866. doi:10.1161/CIRCULATIONAHA.122.059863

28. Zheng X, Yang Y, Huang Fu C, Huang R. Identification and verification of promising diagnostic biomarkers in patients with hypertrophic cardiomyopathy associate with immune cell infiltration characteristics. Life Sci. 2021;285:119956. doi:10.1016/j.lfs.2021.119956

29. Zhao G, Jeoung NH, Burgess SC, Rosaaen-Stowe KA, Inagaki T, Latif S, Shelton JM, McAnally J, Bassel-Duby R, Harris RA, Richardson JA, Kliewer SA. Overexpression of pyruvate dehydrogenase kinase 4 in heart perturbs metabolism and exacerbates calcineurin-induced cardiomyopathy. Am J Physiol Heart Circ Physiol. 2008;294(2):H936–43. doi:10.1152/ajpheart.00870.2007

30. Zannas AS, Jia M, Hafner K, Baumert J, Wiechmann T, Pape JC, Arloth J, Ködel M, Martinelli S, Roitman M, Röh S, Haehle A, Emeny RT, Iurato S, Carrillo-Roa T, Lahti J, Räikkönen K, Eriksson JG, Drake AJ, Waldenberger M, Wahl S, Kunze S, Lucae S, Bradley B, Gieger C, Hausch F, Smith AK, Ressler KJ, Müller-Myhsok B, Ladwig KH, Rein T, Gassen NC, Binder EB. Epigenetic upregulation of FKBP5 by aging and stress contributes to NF-κB-driven inflammation and cardiovascular risk. Proc Natl Acad Sci U S A. 2019;116(23):11370–11379. doi:10.1073/pnas.1816847116

31. Riechert E, Kmietczyk V, Stein F, Schwarzl T, Sekaran T, Jürgensen L, Kamuf-Schenk V, Varma E, Hofmann C, Rettel M, Gür K, Ölschläger J, Kühl F, Martin J, Ramirez-Pedraza M, Fernandez M, Doroudgar S, Méndez R, Katus HA, Hentze MW, Völkers M. Identification of dynamic RNA-binding proteins uncovers a Cpeb4-controlled regulatory cascade during pathological cell growth of cardiomyocytes. Cell Rep. 2021;35(6):109100. doi:10.1016/j.celrep.2021.109100

32. Travaglini KJ, Nabhan AN, Penland L, Sinha R, Gillich A, Sit R V, Chang S, Conley SD, Mori Y, Seita J, Berry GJ, Shrager JB, Metzger RJ, Kuo CS, Neff N, Weissman IL, Quake SR, Krasnow MA. A molecular cell atlas of the human lung from single-cell RNA sequencing. Nature. 2020;587(7835):619-625. doi:10.1038/s41586-020-2922-4

33. Xu P, Yi Y, Xiong L, Luo Y, Xie C, Luo D, Zeng Z, Liu A. Oncostatin M/Oncostatin M Receptor Signal Induces Radiation-Induced Heart Fibrosis by Regulating SMAD4 in Fibroblast. Int J Radiat Oncol Biol Phys. Published online August 2023. doi:10.1016/j.ijrobp.2023.07.033

34. Chen PW, Jian X, Heissler SM, Le K, Luo R, Jenkins LM, Nagy A, Moss J, Sellers JR, Randazzo PA. The Arf GTPase-activating Protein, ASAP1, Binds Nonmuscle Myosin 2A to Control Remodeling of the Actomyosin Network. J Biol Chem. 2016;291(14):7517-7526. doi:10.1074/jbc.M115.701292

35. Manara MC, Pasello M, Scotlandi K. CD99: A Cell Surface Protein with an Oncojanus Role in Tumors. Genes (Basel). 2018;9(3). doi:10.3390/genes9030159

36. Nguyen XX, Muhammad L, Nietert PJ, Feghali-Bostwick C. IGFBP-5 Promotes Fibrosis via Increasing Its Own Expression and That of Other Pro-fibrotic Mediators. Front Endocrinol (Lausanne). 2018;9:601. doi:10.3389/fendo.2018.00601

37. Enomoto Y, Matsushima S, Shibata K, Aoshima Y, Yagi H, Meguro S, Kawasaki H, Kosugi I, Fujisawa T, Enomoto N, Inui N, Nakamura Y, Suda T, Iwashita T. LTBP2 is secreted from lung myofibroblasts and is a potential biomarker for idiopathic pulmonary fibrosis. Clin Sci (Lond). 2018;132(14):1565–1580. doi:10.1042/CS20180435

38. Litviňuková M, Talavera-López C, Maatz H, Reichart D, Worth CL, Lindberg EL, Kanda M, Polanski K, Heinig M, Lee M, Nadelmann ER, Roberts K, Tuck L, Fasouli ES, DeLaughter DM, McDonough B, Wakimoto H, Gorham JM, Samari S, Mahbubani KT, Saeb-Parsy K, Patone G, Boyle JJ, Zhang H, Zhang H, Viveiros A, Oudit GY, Bayraktar OA, Seidman JG, Seidman CE, Noseda M, Hubner N, Teichmann SA. Cells of the adult human heart. Nature. 2020;588(7838):466-472. doi:10.1038/s41586-020-2797-4

39. Chappell JC, Mouillesseaux KP, Bautch VL. Flt-1 (vascular endothelial growth factor receptor-1) is essential for the vascular endothelial growth factor-Notch feedback loop during angiogenesis. Arterioscler Thromb Vasc Biol. 2013;33(8):1952–1959. doi:10.1161/ATVBAHA.113.301805

40. Reichart D, Lindberg EL, Maatz H, Miranda AMA, Viveiros A, Shvetsov N, Gärtner A, Nadelmann ER, Lee M, Kanemaru K, Ruiz-Orera J, Strohmenger V, DeLaughter DM, Patone G, Zhang H, Woehler A, Lippert C, Kim Y, Adami E, Gorham JM, Barnett SN, Brown K, Buchan RJ, Chowdhury RA, Constantinou C, Cranley J, Felkin LE, Fox H, Ghauri A, Gummert J, Kanda M, Li R, Mach L, McDonough B, Samari S, Shahriaran F, Yapp C, Stanasiuk C, Theotokis PI, Theis FJ, van den Bogaerdt A, Wakimoto H, Ware JS, Worth CL, Barton PJR, Lee YA, Teichmann SA, Milting H, Noseda M, Oudit GY, Heinig M, Seidman JG, Hubner N, Seidman CE. Pathogenic variants damage cell composition and single cell transcription in cardiomyopathies. Science. 2022;377(6606):eabo1984. doi:10.1126/science.abo1984

41. Cain SA, Mularczyk EJ, Singh M, Massam-Wu T, Kielty CM. ADAMTS-10 and -6 differentially regulate cell-cell junctions and focal adhesions. Sci Rep. 2016;6:35956. doi:10.1038/srep35956

42. Eraslan G, Drokhlyansky E, Anand S, Fiskin E, Subramanian A, Slyper M, Wang J, Van Wittenberghe N, Rouhana JM, Waldman J, Ashenberg O, Lek M, Dionne D, Win TS, Cuoco MS, Kuksenko O, Tsankov AM, Branton PA, Marshall JL, Greka A, Getz G, Segrè A V, Aguet F, Rozenblatt-Rosen O, Ardlie KG, Regev A. Single-nucleus cross-tissue molecular reference maps toward understanding disease gene function. Science. 2022;376(6594):eabl4290. doi:10.1126/science.abl4290

43. Bajpai G, Schneider C, Wong N, Bredemeyer A, Hulsmans M, Nahrendorf M, Epelman S, Kreisel D, Liu Y, Itoh A, Shankar TS, Selzman CH, Drakos SG, Lavine KJ. The human heart contains distinct macrophage subsets with divergent origins and functions. Nat Med. 2018;24(8):1234–1245. doi:10.1038/s41591-018-0059-x

44. Yano M, Akazawa H, Oka T, Yabumoto C, Kudo-Sakamoto Y, Kamo T, Shimizu Y, Yagi H, Naito AT, Lee JK, Suzuki J ichi, Sakata Y, Komuro I. Monocyte-derived extracellular Nampt-dependent biosynthesis of NAD(+) protects the heart against pressure overload. Sci Rep. 2015;5:15857. doi:10.1038/srep15857

45. Liu J, Wu J, Wang R, Zhong D, Qiu Y, Wang H, Song Z, Zhu Y. ANKRD22 Drives Rapid Proliferation of Lgr5(+) Cells and Acts as a Promising Therapeutic Target in Gastric Mucosal Injury. Cell Mol Gastroenterol Hepatol. 2021;12(4):1433–1455. doi:10.1016/j.jcmgh.2021.06.020

46. Zhang L, Malik S, Pang J, Wang H, Park KM, Yule DI, Blaxall BC, Smrcka A V. Phospholipase Cε hydrolyzes perinuclear phosphatidylinositol 4-phosphate to regulate cardiac hypertrophy. Cell. 2013;153(1):216–227. doi:10.1016/j.cell.2013.02.047

47. Sciarretta S, Volpe M, Sadoshima J. Mammalian target of rapamycin signaling in cardiac physiology and disease. Circ Res. 2014;114(3):549–564. doi:10.1161/CIRCRESAHA.114.302022

48. Frangogiannis NG, Michael LH, Entman ML. Myofibroblasts in reperfused myocardial infarcts express the embryonic form of smooth muscle myosin heavy chain (SMemb). Cardiovasc Res. 2000;48(1):89–100. doi:10.1016/s0008-6363(00)00158-9

49. Roy C, Slimani A, de Meester C, Amzulescu M, Pasquet A, Vancraeynest D, Beauloye C, Vanoverschelde JL, Gerber BL, Pouleur AC. Associations and prognostic significance of diffuse myocardial fibrosis by cardiovascular magnetic resonance in heart failure with preserved ejection fraction. J Cardiovasc Magn Reson Off J Soc Cardiovasc Magn Reson. 2018;20(1):55. doi:10.1186/s12968-018-0477-4

50. Kawasaki T, Sugihara H. Subendocardial ischemia in hypertrophic cardiomyopathy. J Cardiol. 2014;63(2):89–94. doi:10.1016/j.jjcc.2013.10.005

51. Oka T, Akazawa H, Naito AT, Komuro I. Angiogenesis and cardiac hypertrophy: maintenance of cardiac function and causative roles in heart failure. Circ Res. 2014;114(3):565–571. doi:10.1161/CIRCRESAHA.114.300507

52. Ferrari R, Rizzo P. The Notch pathway: a novel target for myocardial remodelling therapy? Eur Heart J. 2014;35(32):2140–2145. doi:10.1093/eurheartj/ehu244

53. Takada Y, Ye X, Simon S. The integrins. Genome Biol. 2007;8(5):215. doi:10.1186/gb-2007-8-5-215

54. Meagher PB, Lee XA, Lee J, Visram A, Friedberg MK, Connelly KA. Cardiac Fibrosis: Key Role of Integrins in Cardiac Homeostasis and Remodeling. Cells. 2021;10(4). doi:10.3390/cells10040770

